# Genomic hallmarks and therapeutic implications of G0 cell cycle arrest in cancer

**DOI:** 10.1101/2021.11.12.468410

**Authors:** Anna J. Wiecek, Stephen J. Cutty, Daniel Kornai, Mario Parreno-Centeno, Lucie E. Gourmet, Guidantonio Malagoli Tagliazucchi, Daniel H. Jacobson, Ping Zhang, Lingyun Xiong, Gareth L. Bond, Alexis R. Barr, Maria Secrier

## Abstract

Therapy resistance in cancer is often driven by a subpopulation of cells that are temporarily arrested in a non-proliferative G0 state, which is difficult to capture and whose mutational drivers remain largely unknown. We developed methodology to robustly identify this state from transcriptomic signals and characterised its prevalence and genomic constraints in solid primary tumours. We show that G0 arrest preferentially emerges in the context of more stable, less mutated genomes which maintain *TP53* integrity and lack the hallmarks of DNA damage repair deficiency, while presenting increased APOBEC mutagenesis. We employ machine learning to uncover novel genomic dependencies of this process and validate the role of the centrosomal gene *CEP89* as a modulator of proliferation/G0 arrest capacity. Lastly, we demonstrate that G0 arrest underlies unfavourable responses to various therapies exploiting cell cycle, kinase signalling and epigenetic mechanisms in single cell data, and propose a G0 arrest transcriptional signature that is linked with therapeutic resistance and can be used to further study and clinically track this state.

## BACKGROUND

Tumour proliferation is one of the main hallmarks of cancer development(1), and has been extensively studied. While most of the cells within the tumour have a high proliferative capacity, occasionally under stress conditions some cells will become arrested temporarily in the G0 phase of the cell cycle, in a reversible state often referred to as ‘quiescence’, ‘dormancy’, ‘diapause-like’, or a potentially irreversible state called ‘senescence’, where they maintain minimal basal activity(2, 3, 4, 5). It has been proposed that G0 arrest enables cells to become resistant to anti-cancer compounds that target actively dividing cells, such as chemotherapy(5, 6, 7). Moreover, a drug-tolerant ‘persister’ cell state represented by slow cycling, entirely quiescent or even senescent cells(4, 8, 9, 10, 11) has been observed in a variety of pre-existing or acquired resistance scenarios, also in the context of targeted therapies(12, 13). As neoplastic cells evolve, G0 arrest can also be employed as a mechanism to facilitate immune evasion(14, 15) or adaptation to new environmental niches during metastatic seeding(16, 17). In the context of disseminated tumour cells, these G0 cycle arrest states can facilitate minimal residual disease, a major cause of relapse in the clinic(18).

Although G0 arrest is a widely conserved cellular state, essential for the normal development and homeostasis of eukaryotes(2, 19), and has been extensively studied in a variety of organisms including bacteria and yeast(20, 21), its role and different facets in cancer are still poorly defined. Hampering our understanding is the fact that it represents a number of heterogeneous states(19, 22). Canonically, cells can be forced into G0 arrest through serum starvation, mitogen withdrawal or contact inhibition(19). Cells can also undergo G0 arrest spontaneously in response to cell-intrinsic factors like replication stress(23, 24, 25). This process is controlled by p53(26), which triggers the inhibition of cyclin-CDK complexes by activating p21(24). This in turn allows the assembly of the DREAM complex - a key effector responsible for repression of cell-cycle dependent gene expression(27). Min and Spencer(28) recently demonstrated a much broader systemic coordination of 198 genes underlying distinct types of G0 arrest by profiling the transcriptomes of cells that entered this state either spontaneously or upon different stimuli. Additionally, proliferation-G0 decisions can be impacted by oncogenic changes such as *MYC* amplification(29) or altered p38/ERK signalling(30).

Despite these advances, the identification of G0-arrested cells within tumours presents an ongoing challenge due to their scarcity and lack of universal, easily measurable markers for the activation and maintenance of this state. As they are often defined by a lack of proliferative markers(31, 32), different forms of G0 arrest such as quiescence, senescence, dormancy and (to a lesser extent) stemness, might sometimes be used interchangeably (4, 33). Quiescent and dormant cells can readily resume their proliferative state, senescent cells are irreversibly arrested(28) while cancer stem cells have a high capacity for self-renewal and sit at the top of the differentiation hierarchy(34). Even though the same cell cycle arrest programme underlies all of these states, they are linked with distinct environmental stimuli, and drive cancer progression and therapeutic resistance in different ways (11, 12, 35, 36). Increasing evidence from the literature points towards the rapid adaptation of tumour cells to drug treatments being enabled by a slow dividing or a quiescent state that persists for a short period of time before the cells start reproliferating(37). Thus, quiescence could more frequently be encountered at the early stages of therapeutic resistance compared to other cell cycle arrest phenotypes, although senescence and stemness are also often discussed in this context. Biomarkers of cell cycle arrest and persistence that are sufficiently specific and robust to be clinically useful are clearly needed.

Furthermore, our understanding of how cancer evolution is shaped by proliferation and G0 arrest decisions is limited. The proliferative heterogeneity of cancer cell populations has been previously described and linked with FAK/AKT1 signalling(38), but the constraints and consequences of these cell state switches have not been systematically profiled across cancer tissues. The extent to which G0 arrest in cancer is enacted through transcriptional or genetic control is unknown(5, 39), and neither are the mutational processes and genomic events modulating this state. Understanding the evolutionary triggers and molecular mechanisms that enable cancer cells to enter and maintain G0 arrest would enable us to develop pharmacological strategies to selectively eradicate these arrested cancer cells or prevent them from re-entering proliferative cycles.

To address these challenges, we have developed a new method to reliably quantify G0 arrest in cancer using transcriptomic data, and employed it to characterise this phenomenon in bulk and single cell datasets from a variety of solid tumours. We describe the spectrum of proliferation and G0 arrest decisions in primary tumours, which reflects a range of stress adaptation mechanisms during the course of cancer development from early to advanced disease. We identify and validate mutational constraints for the emergence of G0 arrest, hinting at potential new therapeutic targets that could exploit this mechanism. We also demonstrate the relevance of G0 arrest to responses to a range of compounds targeting cell cycle, kinase signalling and epigenetic mechanisms in single cell datasets, and propose an expression signature that could be employed to detect treatment resistance induced by G0 arrested tumour cells.

## RESULTS

### Evaluating G0 arrest in cancer from transcriptomic data

We hypothesised that primary tumours contain varying numbers of cells temporarily or permanently arrested in the cell cycle, which reflect evolutionary adaptations to cellular stress and may determine their ability to overcome antiproliferative therapies. To capture this elusive phenotype, we developed a computational framework that would allow us to quantify G0 arrest signals in bulk and single cell sequenced cancer samples (Figure 1a). To define a signature of G0 arrest, we focused on genes that have been shown by Min and Spencer(28) to be specifically activated or inactivated during quiescence that arises spontaneously or as a response to serum starvation, contact inhibition, MEK inhibition or CDK4/6 inhibition. The activity of 139 of these genes changed in a coordinated manner across all these five distinct forms of quiescence, likely representing generic transcriptional consequences of G0 arrest. The expression levels of these markers were used to derive a score reflecting the relative abundance of G0 arrested cells within individual tumours (see Methods, Supplementary Table 1).

**Figure 1:**
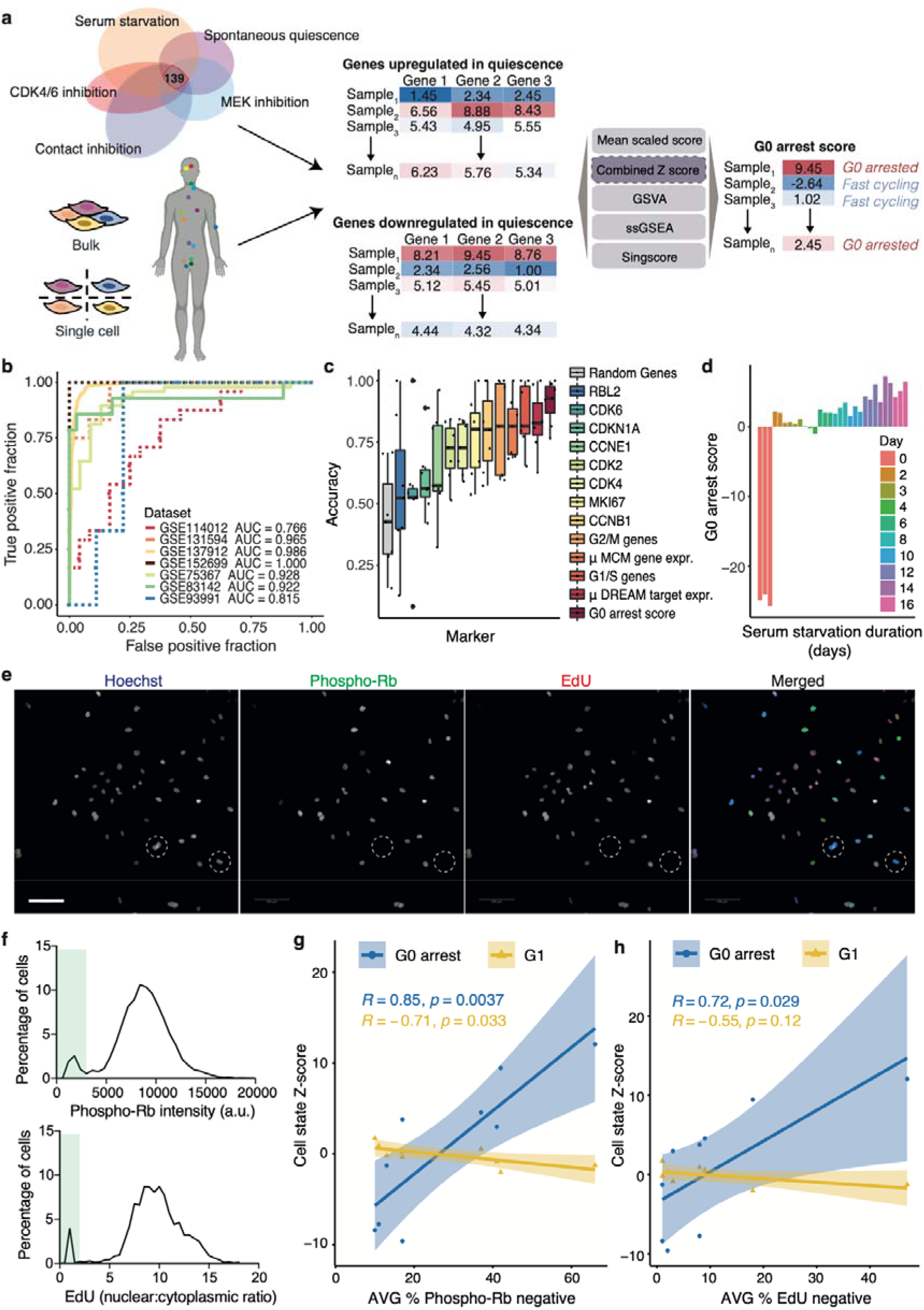
Methodology for quantifying G0 arrest in cancer. **(a)** Workflow for evaluating G0 arrest from RNA-seq data. 139 genes differentially expressed in multiple forms of quiescence were employed to score G0 arrest across cancer tissues. **(b)** Receiver operating characteristic (ROC) curves illustrating the performance of the z-score methodology on separating actively proliferating and G0 arrested cells in seven single-cell (continuous curves) and bulk RNA-seq (dotted curves) datasets. AUC = area under the curve. **(c)** Compared classification accuracies of the G0 arrest z-score approach and classic cell proliferation markers across the seven single-cell/bulk RNA-seq validation datasets. **(d)** G0 arrest levels of embryonic fibroblast cells under serum starvation for various amounts of time. Replicates are depicted in the same colour. **(e)** Representative images of lung cancer cell lines immunostained and analysed to detect the G0 arrest fraction. Hoechst (labels all nuclei) is in blue, phospho-Rb in green and EdU in red in merged image. White dashed circles highlight G0 arrested cells that are negative for both phospho-Rb and EdU signals. Scale bar: 100 µm. **(f)** Graphs show single cell quantification of phospho-Rb and EdU intensities taken from images and used to define the cut-off to calculate the G0 arrest fraction (green boxes). Images in **(e)** and graphs in **(f)** are taken from the A549 cell line. **(g-h)** Correlation between theoretical estimates of a G0 or G1 state and the fraction of cells entering G0 arrest in nine lung adenocarcinoma cell lines, as assessed through **(g)** phospho-Rb assays and **(h)** EdU assays. Mean of n=3 is shown for the average percentage of G0 arrested cells.

To validate this signature and select the optimal method to score G0 arrest in individual samples amongst different enrichment/rank-based scoring methodologies(40, 41, 42, 43), we used seven single-cell and bulk datasets(12, 44, 45, 46, 47, 48, 49) where actively proliferating and quiescent/dormant cells had been independently isolated and sequenced (Supplementary Table 2, Methods). We tested the performance of our signature and scoring methodology, as well as that of other commonly used gene signatures, in distinguishing between the truly quiescent/dormant and truly proliferating cells in these seven datasets while varying the expression cut-offs for labelling cells as G0 arrested or proliferating based on the respective signature. A combined Z-score approach had the highest accuracy in detecting signals of G0 arrest, with a 91% mean performance in classifying cells as G0 arrested or cycling (Figure 1b, Supplementary Figure 1a-b). Indeed, the individual cells that had been identified as arrested in G0 in the experiments showed a significantly higher Z-score than the dividing cells across all datasets (Supplementary Figure 1c). Our signature reflected an expected increase in p27 protein levels, which are elevated in during G0 arrest(50) (Supplementary Figure 1d). It also outperformed classical cell cycle and arrest markers, such as the expression of targets of the DREAM complex, CDK2, Ki67 and of mini-chromosome replication maintenance (MCM) protein complex genes - which are involved in the initiation of eukaryotic genome replication, as well as recently defined G1/S and G2/M signatures (51) (Figure 1c). Importantly, our approach provided a good separation between G0 and proliferating samples across a variety of cancer types and models including cancer cell lines, 3D organoid cultures, circulating tumour cells and patient-derived xenografts (Supplementary Table 2), thereby demonstrating its broad applicability. Furthermore, the strength of the score appeared to reflect the duration of G0 arrest (52) (Figure 1d).

We further experimentally validated our methodology in nine lung adenocarcinoma cell lines. We estimated the fraction of G0 cells in each of these cell lines using quantitative, single-cell imaging of phospho-Ser807/811-Rb (phospho-Rb, which labels proliferative cells(53)) and 24 hour EdU proliferation assays (Figure 1e-h). In these assays, cells were pulse-labelled with the nucleotide analogue, EdU, for 24h before fixation and immunostaining. Only cells which have proliferated in the last 24h will be labelled by EdU. EdU negative cells are classed as G0. Cells that were negative for phospho-Rb were also defined as G0, and not G1, since they have not yet passed the restriction point (phospho-Rb negative; see Methods, Figure 1e-f). This G0 fraction was further validated in A549 and NCI-H1944 cells where endogenous PCNA has been labelled with an mRuby fluorophore to enable tracking of cell cycle phase lengths by live-cell imaging (Zerjatke et al(54), Methods). By quantifying the G0/G1 length in individual cells (i.e. time taken to enter S-phase after mitotic exit) over a 48h period, we could see that these cells were quiescent and not senescent (or in deep quiescence), as all G0/G1 cells did eventually enter S-phase, albeit with variable timing (Supplementary Figure 1e).

There was a remarkably good correlation between our predicted G0 arrest levels based on the expression of these cell lines from the Cancer Cell Line Encyclopedia (CCLE) and the fraction of G0 cells in the experiment as assessed by lack of EdU incorporation over a 24h period (EdU incorporation only occurs during S phase) but particularly by lack of Rb phosphorylation. Phosphorylation and inactivation of the retinoblastoma protein is often used to define the boundary between G0 and G1, and was specifically shown to distinguish the G0 state recently by Stallaert et al(53). Furthermore, a G1 signature (Methods) was not associated with these experimental measurements, suggesting our method recovers a state more similar to G0 arrest rather than a prolonged G1 state (Figure 1g-h). The G0 arrest correlations appeared robust to random removal of individual genes from the signature, with no single gene having an inordinate impact on the score (Supplementary Figures 1f-h). This provided further reassurance that our Z-score based methodology is successful in capturing G0 arrest signals from bulk tumour data.

### The spectrum of G0 arrest capacity in solid primary tumours

Having established a robust framework for quantifying G0 cell cycle arrest in cancer, we next profiled 8,005 primary tumour samples across 31 solid cancer tissues from The Cancer Genome Atlas (TCGA). After accounting for potential confounding signals of non-cycling non-tumour cells from the microenvironment by correcting for tumour purity (see Methods, Supplementary Figures 1i-j), we observed an entire spectrum of fast proliferating to slowly cycling tumours, with the latter presenting stronger G0-linked signals (Figure 2a). While we acknowledge that no tumour would be entirely quiescent/senescent and we cannot identify individual G0 arrested cells within the tumour, this analysis does capture a broad range of phenotypes reflecting varying proliferation and cell cycle arrest rates, which suggests that G0 arrest is employed to different extents by tumours as an adaptive mechanism to various extrinsic and intrinsic stress factors. Cancers known to be frequently dormant, such as glioblastoma(6, 44), were amongst the highest ranked in terms of G0 arrest levels, along with kidney and adrenocortical carcinomas (Figure 2b). This is likely explained by the innate proliferative capacity of the respective tissues. Indeed, tissues with lower stem cell division rates presented a greater propensity for G0 arrest (Figure 2c)(55).

**Figure 2:**
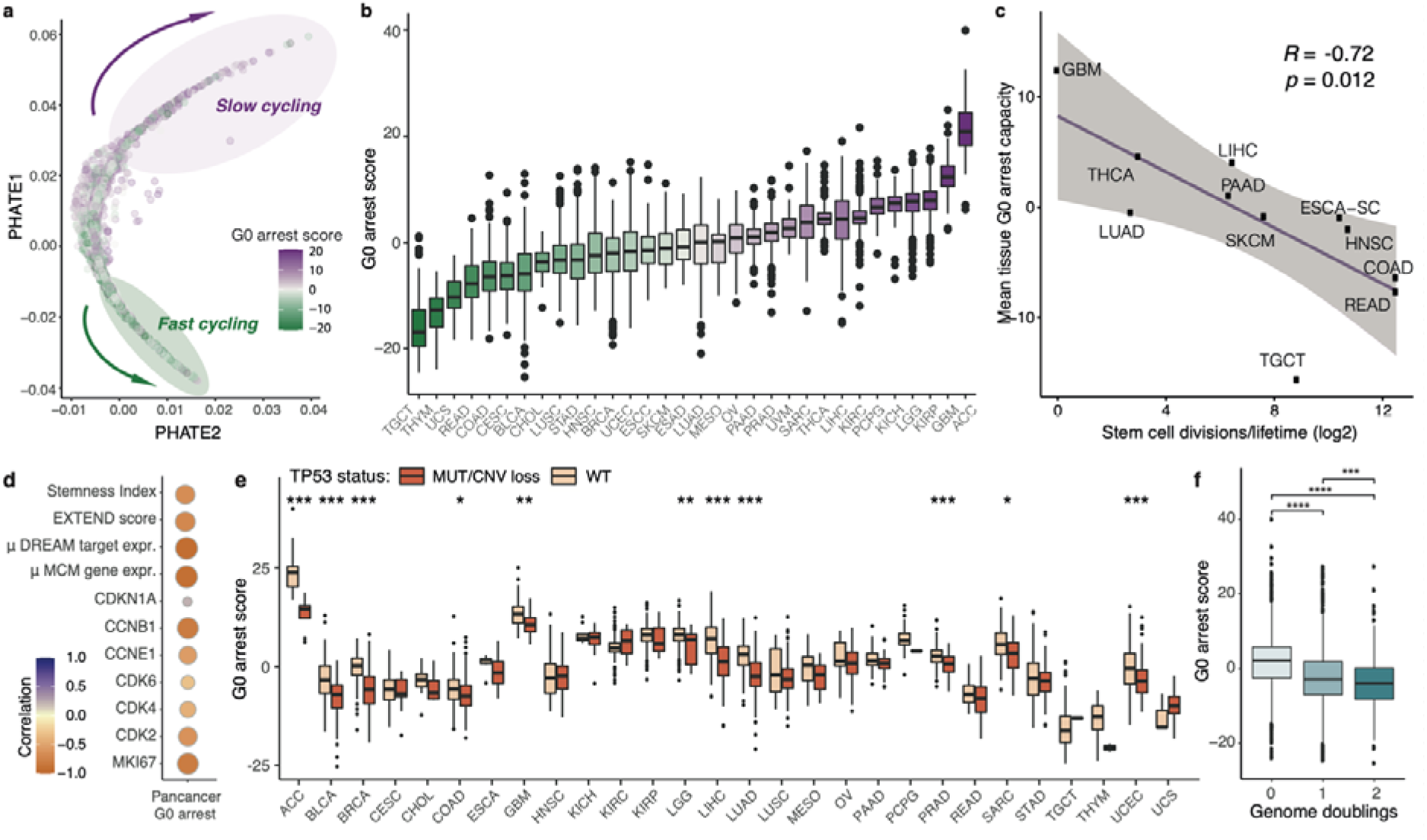
Pan-cancer evaluation of proliferative heterogeneity and linked tumour hallmarks. **(a)** PHATE plot illustrating the wide spectrum of proliferative to slow cycling/arrested states across 8,005 primary tumour samples from TCGA. Each sample is coloured according to the relative G0 arrest level. **(b)** Variation in tumour G0 arrest levels across different cancer tissues. **(c)** Correlation between mean G0 arrest capacity and stem cell division estimates for various tissue types. **(d)** Correlating tumour G0 arrest scores with cancer cell stemness (Stemness Index), telomerase activity (EXTEND score), p21 activity (CDKN1A) and the expression of several commonly used proliferation markers. The Pearson correlation coefficient is displayed. RC – replication complex. **(e)** Consistently higher levels of G0 arrest are detected in samples with functional p53. **(f)** Lower G0 arrest scores are observed in tumours with one or two whole genome duplication events. Wilcoxon rank-sum test p-values are displayed in boxplots, *p<0.05; **p<0.01; ***p<0.001; ****p<0.0001.

Our score showed strong negative correlations with the expression of proliferation markers (Figure 2d), suggesting that it captures a cellular state that could potentially act as a baseline for all major forms of cell cycle arrest, including quiescence, senescence, stemness and clinical dormancy. Indeed, we found that our signature could to a certain extent also separate senescent cells from proliferating ones in single cell data from Hernandez-Segura et al(56), which is unsurprising since these cells are also in the G0 phase (Supplementary Figure 2a). Indeed, the authors of this resource highlight that some of the pathways uncovered in these senescent cells may be shared with quiescence, which is also backed up by a study from Fujimaki and Yao(57) suggesting similarities between deep quiescence and senescence. Our score did not show strong correlations with other markers of senescence such as the Senescence-Associated Secretory Phenotype (SASP) and β- galactosidase activity(58, 59, 60) in single cells or TCGA samples (Figure 2d, Supplementary Figures 2b-d), although we cannot exclude the possibility of a senescent state being captured occasionally given that neither β-galactosidase nor the SASP are obligatory for maintaining senescence(61). However, the underpinning programme appears to be distinct from that of cancer stem cells, marked by signatures associated with high telomerase activity and an undifferentiated state(62, 63) (Supplementary Figure 2e-f).

Lastly, we confirmed expected dependencies on the p53/p21/DREAM activation axis: tumours that were proficient in *TP53* or the components of the DREAM complex, as well as those with higher p21 expression, had elevated G0 arrest levels across numerous tissues (Figure 2e, Supplementary Figures 2g-h), although only 8 out of 139 genes in our signature are directly transcriptionally regulated by p53(64). Nevertheless, p53 proficiency appears to be a non-obligatory dependency of G0 arrest, which is also observed to arise in p53 mutant scenarios in 21% of cases. p53 has also been shown to play a role in preventing the occurrence of larger structural events and polyploidy(65, 66, 67), potentially explaining the lower G0 arrest levels we observed in tumours that had undergone whole genome duplication (Figure 2f).

### The genomic background of G0 arrest in cancer

Cancer evolution is often driven by a variety of genomic events, ranging from single base substitutions to larger scale copy number variation and rearrangements of genomic segments. It is reasonable to expect that certain mutations accumulated by the cancer cells might enable a more proliferative phenotype, impairing the ability of cells to enter G0 arrest, or – on the contrary – might favour cell cycle exit as a temporary adaptive mechanism to extreme levels of stress. Having obtained G0 arrest estimates for primary tumour samples, we set out to identify potential genomic triggers or constraints that may shape proliferation versus G0 arrest decisions in cancer. We identified 285 cancer driver genes that were preferentially altered (via mutations or copy number alterations) either in slow cycling or fast proliferating tumours (Figure 3a). Reassuringly, this list included genes previously implicated in driving cell cycle exit decisions such as *TP53* and *MYC*(26, 29). We also investigated associations with mutagenic footprints of carcinogens (termed “mutational signatures”), which can be identified as trinucleotide substitution patterns in the genome(68, 69). 15 mutational signatures were linked with G0 arrest levels either within individual cancer studies or pan-cancer (Supplementary Figure 2i).

**Figure 3:**
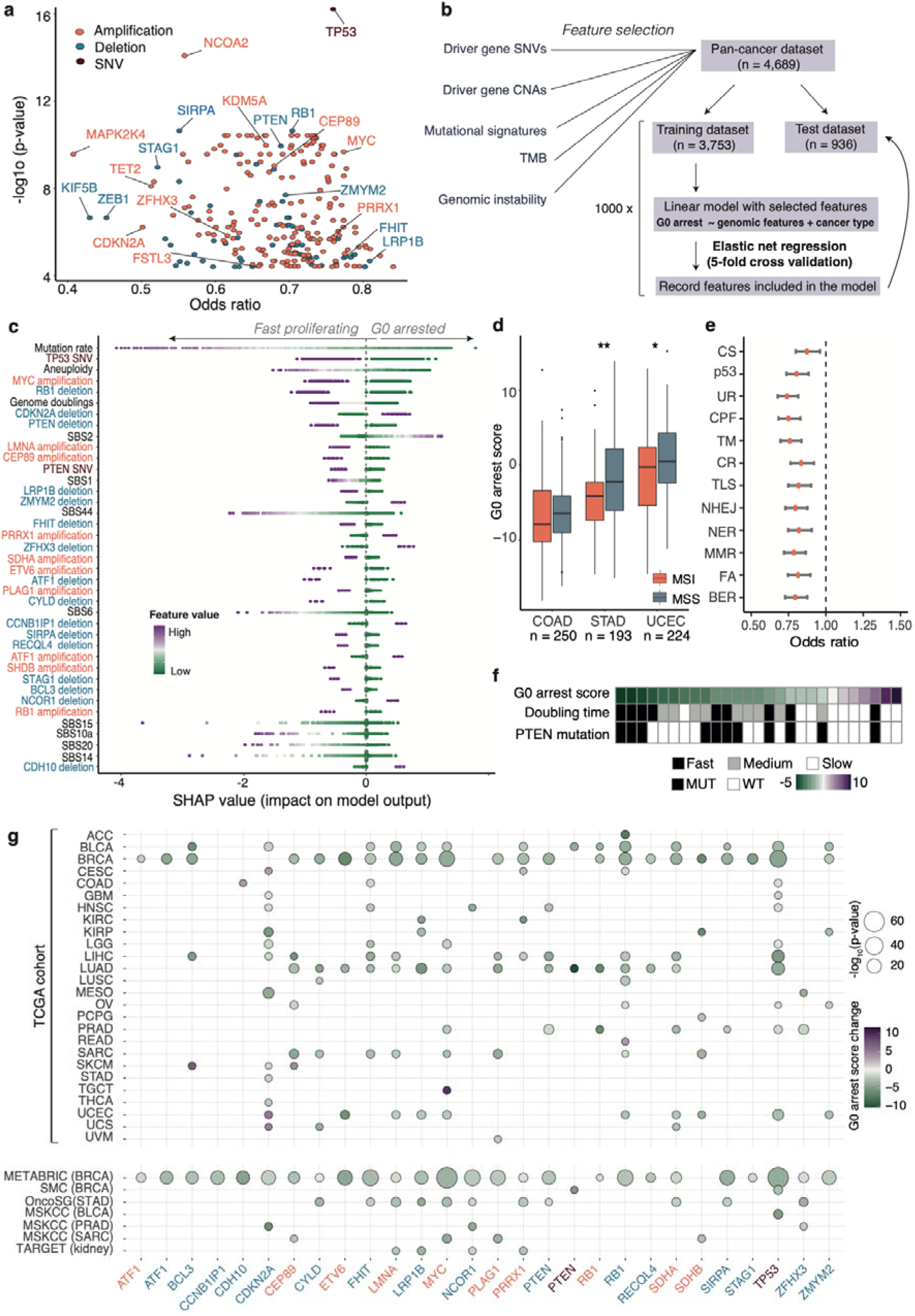
Genomic landscape of G0 arrest decisions in cancer. **(a)** Cancer drivers with mutations or copy number alterations depleted pan-cancer in a G0 arrest context. Features further selected by the pan-cancer model are highlighted. **(b)** Schematic of the ensemble elastic net modelling employed to prioritise genomic changes associated with G0 arrest. **(c)** Genomic events significantly associated with G0 arrest, ranked according to their importance in the model (highest to lowest). Each point depicts an individual tumour sample, coloured by the value of the respective feature. For discrete variables purple indicates the presence of the feature and green its absence. The Shapley values indicate the impact of individual feature values on the G0 arrest score prediction. **(d)** G0 arrest levels are significantly reduced in microsatellite unstable (MSI) samples in stomach adenocarcinoma (STAD) and uterine corpus endometrial carcinoma (UCEC), with the same trend (albeit not significant) shown in colon adenocarcinoma (COAD). Wilcoxon rank-sum test *p<0.05; **p<0.01. **(e)** Genomic alterations are depleted across DNA repair pathways during G0 arrest. Odds ratios of mutational load on pathway in G0 arrest are depicted, along with confidence intervals. CS=chromosome segregation; p53=p53 pathway; UR=ubiquitylation response; CPF=checkpoint factors; TM=telomere maintenance; CR=chromatin remodelling; TLS=translesion synthesis; NHEJ=non-homologous end joining; NER=nucleotide excision repair; MMR=mismatch repair; FA=Fanconi Anaemia; BER=base excision repair. **(f)** G0 arrest scores are increased in cell lines with slow doubling time across MCF7 strains, which also show lower prevalence of PTEN mutations. **(g)** Tissue-specific changes in G0 arrest between samples with/without quiescence-associated deletions (blue), amplifications (red) and SNVs (brown) within the TCGA cohort (top) and external validation datasets (bottom).

Following the initial prioritisation of putative genomic constraints of G0 arrest, we employed machine learning to identify those events that could best distinguish slow cycling tumours with higher abundance of G0 arrested cells from fast proliferating ones, while accounting for tissue effects. An ensemble elastic net selection approach similar to the one described by Pich et al(70) was applied for this purpose (Figure 3b, Methods). Our pan-cancer model identified tissue type to be a major determinant of G0 arrest levels (Supplementary Figure 3a). It also uncovered a reduced set of 57 genomic events linked with proliferation/G0 arrest switches, including SNVs and copy number losses in 17 cancer genes, as well as amplifications of 10 cancer genes (Figure 3c). These events could then be successfully employed to predict G0 arrest in a separate test dataset, thus internally validating our model (Supplementary Figure 3b). Thus, while these events are not necessarily causative, the link is strong enough to be identifying G0 arrest states from genomic data alone. Such events may also pinpoint cellular vulnerabilities that could be exploited therapeutically.

Overall, the genomic dependencies of G0 arrest mainly comprised genes involved in cell cycle pathways, p53 regulation and ubiquitination (most likely of cell cycle targets), and RUNX3 regulation, which have previously been shown to play a role in controlling proliferation and cell cycle entry(71) (Supplementary Figure 3c). Invariably, this analysis has captured several events that are well known to promote cellular proliferation in cancer: this is expected and confirms the validity of our model. It was reassuring that a functional *TP53*, lack of *MYC* amplification and lower mutation rates (Figure 3c) were amongst the top ranked characteristics of tumours with high levels of G0 arrest, which also displayed less aneuploidy. However, our analysis has also uncovered novel dependencies of G0 arrest-proliferation decisions that have not been reported previously, such as *CEP89* and *LMNA* amplifications observed in fast cycling tumours, or *ZMYM2* deletions prevalent in samples with high levels of G0 arrest. ZMYM2 has recently been described as a novel binding partner of B-MYB and has been shown to be important in facilitating the G1/S cell cycle transition(72). p16 (*CDKN2A*) deletions, one of the frequent early events during cancer evolution(73, 74), were enriched in tumours with high proportions of cells in G0. *RB1* deletions and amplifications were both associated with a reduction in G0 arrest, which might reflect the dual role of RB1 in regulating proliferation and apoptosis(75).

Our model also calls to attention to the broader mutational processes associated with this cellular state. Such processes showed fairly weak and heterogeneous correlations with G0 arrest within individual cancer tissues (Supplementary Figure 2g), but their contribution becomes substantially clearer pan-cancer once other genomic sources are accounted for. In particular, we identified an association between G0 arrest and mutagenesis induced by the AID/APOBEC family of cytosine deaminases as denoted by signature SBS2(68) (Figure 3c). As highlighted by Mas-Ponte and Supek(76), APOBEC/AID driven mutations tend to be directed towards early-replicating, gene-rich regions of the genome, inducing deleterious events on several genes including *ZMYM2*, which our pan-cancer model has linked with G0 arrest.

In turn, defective DNA mismatch repair, as evidenced by signatures SBS44, SBS20, SBS15, SBS14 and SBS6(68), was prevalent in fast cycling tumours (Figure 3c). Mismatch repair deficiencies lead to hypermutation in a phenomenon termed “microsatellite instability” (MSI), which has been linked with increased immune evasion(77). Cancers particularly prone to MSI include colon, stomach and endometrial carcinomas(78), where this state was indeed linked with reduced G0 arrest (Figure 3d). Furthermore, tumours with high proportions of cells in G0 were depleted of alterations across all DNA damage repair pathways (Figure 3e).

Our measurements of G0 arrest also reflected expected cycling patterns across 27 MCF7 strains(79): cell lines with longer doubling times exhibited increased G0 arrest (Figure 3f). This coincided with a depletion of *PTEN* mutations, a dependency highlighted by the pan-cancer model.

When checking for dependencies in individual cancer tissues, 24 out of the 25 genes identified by the model were significantly associated with G0 arrest or proliferation decisions in at least one tissue, most prominently in breast, lung and liver cancers which also represent the largest studies within TCGA (Figure 3g, top panel). Most of these genomic insults were linked with a decrease in G0 arrest. In external validation datasets these associations, including deletions in *PTEN* and *LRP1B* or amplifications of *MYC*, *CEP89* and *ETV6*, featured most prominently in the largest cohort of breast cancer samples (Figure 3g, bottom panel). These results highlight the fact that although a pan-cancer approach is suited to capture genomic events that are universally associated with cell cycle exit, certain genetic alterations may facilitate a higher or lower propensity of G0 arrest in a single tissue only.

Indeed, when building a tissue-specific breast cancer model of G0 arrest using a combined ANOVA and random forest classification approach (Supplementary Figure 4a), we not only recovered the associations with the *TP53, MYC*, *LMNA* and *ETV6* events already seen in the pan-cancer model (Supplementary Figure 4b), but identified additional events which validated in the METABRIC cohort and were also seen in several other cancers, e.g. bladder, lung and lower grade glioma (Supplementary Figure 4c). Notably, the APOBEC mutational signature SBS2 was the strongest genomic signal linked with G0 arrest in breast cancer (Supplementary Figures 4b,d) and was most prevalent in Her2+ tumours, although the Luminal A subtype showed the highest levels of G0 arrest overall, as expected given its well-known lower proliferative capacity(80) (Supplementary Figures 4e-f).

**Figure 4:**
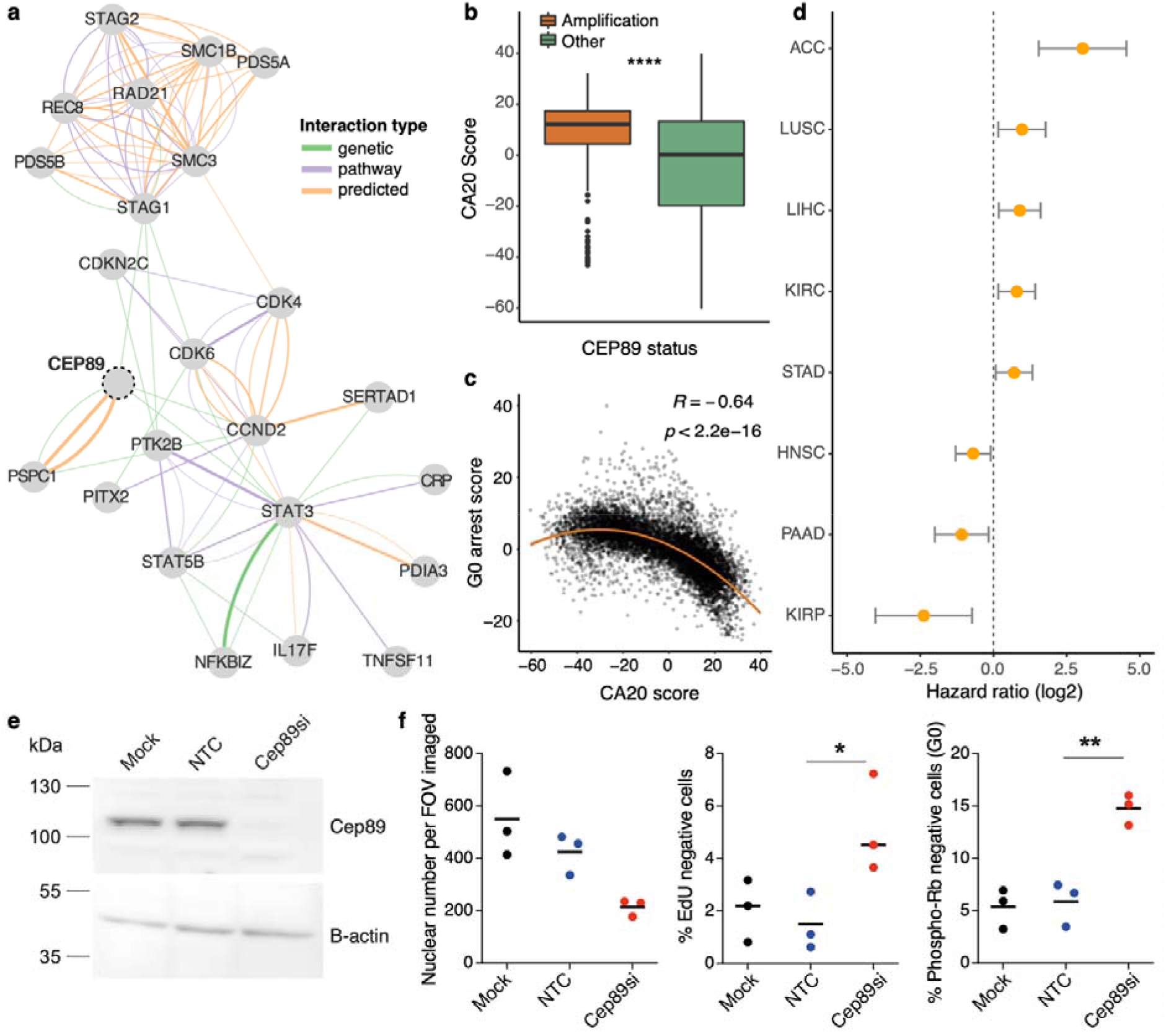
CEP89 amplification is associated with lower G0 arrest capacity. **(a)** Network illustrating CEP89 interactions with cell cycle genes (from GeneMania). The edge colour indicates the interaction type, with green representing genetic interactions, orange representing predicted interactions and purple indicating pathway interactions. The edge width illustrates the interaction weight. **(b)** CA20 scores are significantly increased in TCGA primary tumours containing a *CEP89* amplification. **(c)** Pan-cancer relationship between CA20 and G0 arrest scores across the TCGA cohort. **(d)** Cox proportional hazards analysis estimates of the log hazards ratio for the impact of *CEP89* expression on patient prognosis within individual cancer studies, after adjusting for tumour stage. Patients with high expression of *CEP89* show significantly worse prognosis within ACC, LUSC, LIHC, KIRC and STAD, but significantly better prognosis within HNSC, PAAD and KIRP studies. **(e)** Western blot showing depletion of Cep89 protein 48h after siRNA transfection of NCI- H1299 cells. Mock is lipofectamine only, NTC is Non-targeting control siRNA. B-actin is used as a loading control. **(f)** Graphs show that Cep89 depletion in NCI-H1299 cells leads to a reduction in nuclear number and an increase in the fraction of G0 arrested cells, measured by an increase in the percentage of EdU negative (24h EdU pulse) and Phospho-Ser 807/811 Rb negative cells. One-Way ANOVA, *p<0.05, **p<0.01. Mean of n=3.

### Validation of CEP89 as a modulator of G0 arrest capacity

To gain more insight into the underlying biology of G0 arrest in cancer, we sought to experimentally validate associations highlighted by the pan-cancer model. We focused on the impact of *CEP89* activity on proliferation/arrest decisions due to the high ranking of this putative oncogene in the model, the relatively unexplored links between *CEP89* and cell cycle control, as well as its negative association with G0 arrest across a variety of cancer cell lines (Supplementary Figures 5a-c). The function of *CEP89* is not well characterised, however, the encoded protein has been proposed to function as a centrosomal-associated protein(81, 82). Centrosomes function as major microtubule-organising centres in cells, playing a key role in mitotic spindle assembly(83) and the mitotic entry checkpoint(84). Moreover, centrosomes act as sites of ubiquitin-mediated proteolysis of cell cycle targets(85), and members of several growth signalling pathways, such as Wnt and NF-kB, localise at these structures(86, 87). Several genetic interactions have also been reported between *CEP89* and key cell cycle proteins, including cyclin D2(88) (Figure 4a).

**Figure 5:**
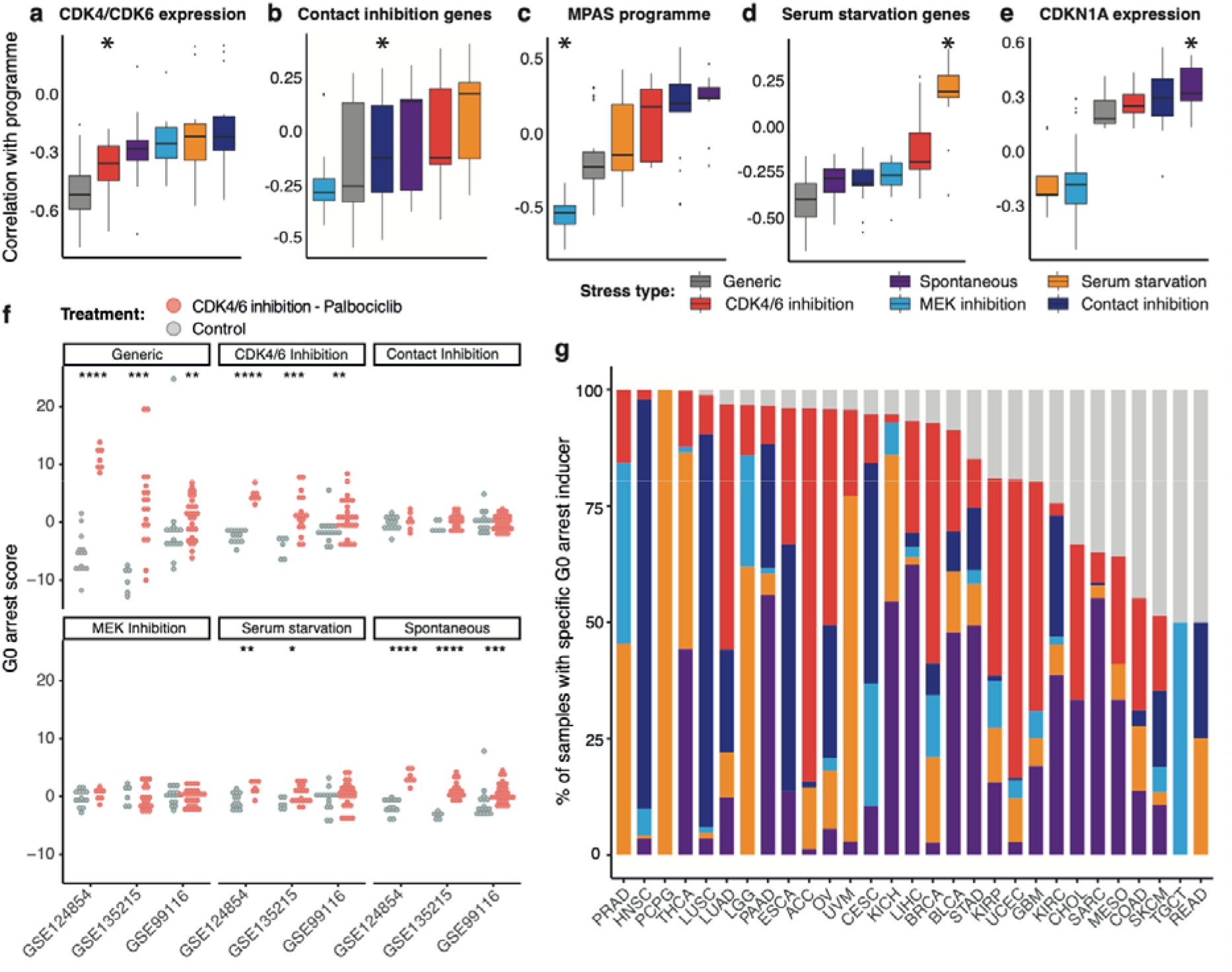
Pan-cancer characterisation of individual G0 stress response programmes. **(a-e)** Comparison of correlation coefficients between stress response programme scores and **(a)** mean expression of CDK4 and CDK6, **(b)** mean expression of curated contact inhibition genes, **(c)** a transcriptional MAPK Pathway Activity Score (MPAS), **(d)** mean expression of curated serum starvation genes, and **(e)** CDKN1A expression, across TCGA cancers. The correlations expected to be strongest (either negative or positive) are denoted by an asterisk. The generic G0 arrest score refers to scores calculated using the original list of 139 genes differentially expressed across all 5 forms of G0 arrest. **(f)** Comparison of stress response programme scores measured in cancer cell lines before (grey) and after (red) Palbociclib treatment across three validation studies. Datasets used for validation are denoted by their corresponding GEO series accession number. **(g)** Predicted stress response diversity in samples with high levels of G0 arrest across individual cancer types. The same colour legend as in **(a)** is applied. Gray bars represent the proportion of samples for which the G0 arrest inducer could not be estimated.

Our model linked *CEP89* amplification with fast cycling tumours (Figure 3c). Centrosome amplification is a common feature of tumours with high proliferation rates and high genomic instability(89), and overexpression of centrosomal proteins can alter centriole structure(90, 91). Indeed, *CEP89* amplified tumours presented elevated expression of a previously reported centrosome amplification signature (CA20)(89) (Figure 4b), which was strongly anticorrelated with G0 arrest levels (Figure 4c). Furthermore, *CEP89* expression was prognostic across multiple cancer tissues (Figure 4d) and linked with toxicity of several cancer compounds in cell line models (Supplementary Figure 5d).

We validated this target in the lung adenocarcinoma cell line NCI-H1299 showing high levels of *CEP89* amplification. Cep89 depletion via siRNA knockdown caused a consistent decrease in cell number, in the absence of any detectable cell death, and an increase in the fraction of G0 cells as measured by phospho-Rb and EdU assays (Figure 4e-f). Thus, we propose *CEP89* as a novel cell proliferation regulator that may be exploited in certain scenarios to control tumour growth.

### Characterisation of individual stress response programmes of G0 arrest

While we had previously examined a generic programme of G0 arrest, cancer cells can enter this state due to different stimuli(19) and this may inform its aetiology and manifestation. To explore this, we re-scored tumours based on gene expression programmes specific to serum starvation, contact inhibition, MEK inhibition, CDK4/6 inhibition or spontaneously occurring quiescence as defined by Min and Spencer(28) (see Methods). We observed a good correlation between our estimates representing individual stress response programmes and the expression of genes associated with the corresponding form of G0 arrest in the literature (Figure 5a-e, Supplementary Figure 6, Methods). Specifically, strong inverse correlations were seen between our CDK4/6 inhibition scores and the mean expression of *CDK4* and *CDK6*, or between our MEK inhibition scores and the expression of genes involved in the MAPK pathway(92). Spontaneous quiescence and serum starvation scores were most correlated with the activity of p21, or of genes involved in the cellular response to starvation, respectively. The contact inhibition programme was also captured, but with lesser specificity. CDK4/6 inhibition-induced G0 arrest levels were further validated using external RNA-seq datasets from cancer cell lines and xenograft mice sequenced before and after treatment with the CDK4/6 inhibitor Palbociclib(93, 94) (Figure 5f, Supplementary Table 3). The fact that the estimates for a generic G0 arrest phenotype were equally or, in some cases, more discriminative of cells treated with Palbociclib confirms the generalisability of this score, which may be more broadly applicable to different tissues and/or model systems, as shown previously in the single cell validation data. The CDK4/6 inhibition scores outperformed all the other stress response subtype scores, suggesting that a combination of individual programmes and the generic score might best identify a specific stimulus driving G0 arrest. Interestingly, we also observed significant differences in spontaneous quiescence scores before and after treatment. Indeed, p21 activity has been linked with the Palbociclib mechanism of action(95, 96), and this analysis suggests potential similarities between CDK4/6 inhibition and p21-dependent G0 arrest phenotypes.

Having validated our framework for quantifying stimulus-specific G0 arrest programmes, we proceeded to estimate the dominant form of stress that may induce cell cycle arrest in different cancer types (Figure 5g). We found a range of G0 arrest aetiologies across most tissues, while a minority of cancers were dominated by a single form of stress response, e.g. serum starvation in all G0 arrested pheochromocytomas and paragangliomas, contact inhibition in 88% of head and neck carcinomas and CDK4/6 inhibition in 80% of adrenocortical carcinomas. While we do not wish to claim that the state of cell cycle arrest will have necessarily been induced by the actual predicted stimulus (impossible in the case of CDK4/6 or MEK inhibition, as the analysed samples are all treatment-naïve), we suggest that the downstream signalling cascade may resemble that triggered by such stimuli, e.g. via CDK4/6 or MEK loss of function mutations.

Some of the differences observed might be explained by the dependency between p53 activity and the form of stress response that is enacted. Amongst the five different forms, spontaneous G0 arrest appeared most strongly dependent on p53 functionality, with a nearly two-fold enrichment of p53 proficient tumours in this group (Supplementary Figure 6f). Indeed, significantly higher levels of spontaneous G0 arrest were observed in the majority of cancers (56%) when p53 was functional rather than mutated. The second most dependent state was that of CDK4/6 inhibition, with increased levels in 36% of cancer types displaying p53 proficiency (Supplementary Figure 6g). Overall, these analyses of stress response states point to common transcriptional features of drug-tolerant G0 arrested cells in different cancer settings that could be employed in designing ways to eradicate these cells in the future.

### Role of G0 arrest in driving therapeutic resistance in cancer uncovered from single cell data

Overall, G0 arrest appears to be beneficial for the long-term outcome of cancer patients, even when accounting for potential confounders such as stage, sex and tissue (Figure 6a, Supplementary Figure 7a). No clear relation was observed between G0 arrest levels within the primary tumour and risk of relapse, although higher G0 arrest was occasionally deemed favourable to avoiding disease recurrence or progression (Supplementary Figure 7b-e). Indeed, such slow cycling, indolent tumours would have higher chances of being eradicated earlier in the disease, which is consistent with reported worse prognosis of patients with higher tumour cell proliferation rates (97). As expected, G0 arrest levels were increased in stage 1 tumours, although later stages also exhibited this phenotype occasionally (Supplementary Figure 7f). However, outcomes do vary depending on the stress source, with worse survival observed upon contact inhibition (Figure 6b). The outcomes also vary by tissue: when the cut-offs between increased G0 arrest and high proliferation were defined on an individual cancer basis rather than pan-cancer, we found that lung, colon or esophageal carcinoma patients displayed significantly worse prognosis in the context of high proportions of G0 arrested cells in the tumour (Figure 6c, Methods). Indeed, p53 wild type colorectal cancers expressing a quiescence-linked fetal phenotype have been recently associated with metastasis and poor prognosis(98). In contrast, adrenocortical and kidney papillary cell carcinoma ranked in the top of cancers with improved survival. It is noticeable that the cancers in the former, worse prognosis group are also amongst the ones displaying lower than average G0 arrest (Figure 2b), so the observed inferior outcomes could in part be linked to these cancers being intrinsically faster progressing. It is possible there is a lower limit below which G0 arrest stops being useful for delaying growth and becomes detrimental instead, perhaps in conjunction with treatment. Indeed, other factors such as the type of therapy received could play a role too. While we are limited in the investigation of such factors in TCGA due to the incomplete records available, these discrepancies should be subject to future research.

**Figure 6:**
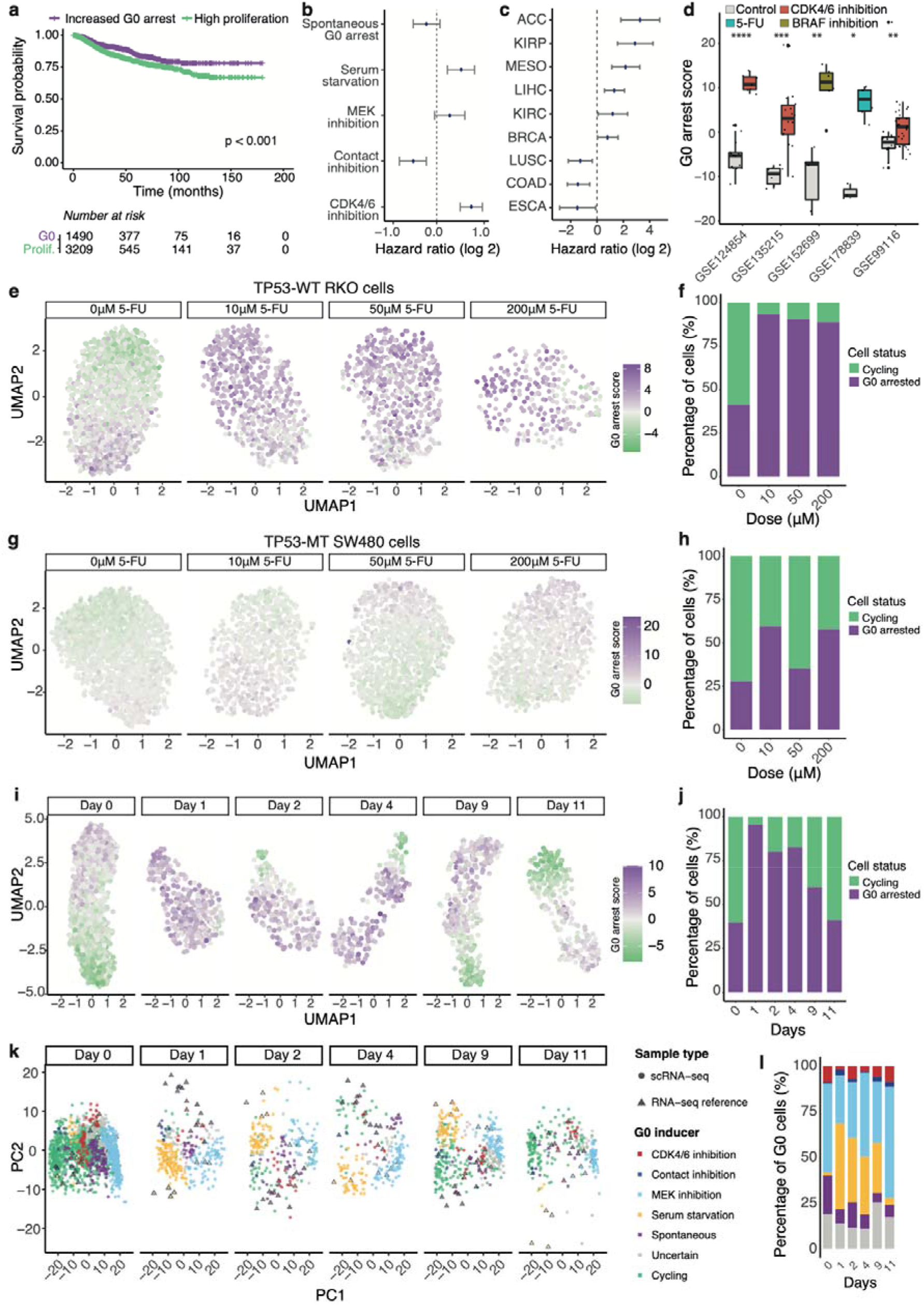
Impact of G0 arrest on patient prognosis and treatment response. **(a)** Disease-specific survival based on proliferation/G0 arrest levels for patients from TCGA within 15 years of follow up. Patients with increased levels of G0 arrest in primary tumours showed significantly better prognosis than patients with fast proliferating tumours. **(b-c)** Hazard ratio ranges illustrating the impact of different forms of G0 induction **(b)** and different tissues **(c)** on patient prognosis, after taking into account potential confounding factors. Values above 0 indicate significantly better prognosis when tumours contain high proportions of cells arrested in G0. **(d)** Change in G0 arrest scores inferred from bulk RNA-seq across breast, pancreatic, colorectal and skin cancer cells in response to treatment with the CDK4/6 inhibitor Palbociclib, 5-FU or the BRAF inhibitor Vemurafenib. **(e-f)** UMAP plot illustrating the response of the TP53-proficient RKO colorectal cancer cell line to various 5-FU doses and the corresponding proportions of cells predicted to be arrested/proliferating. Each dot is an individual cell, coloured according to its G0 arrest level. **(g-h)** The same as previous, but for the TP53-deficient SW480 cell line. **(i-j)** UMAP plot illustrating the response of individual PC9 NSCLC cells to the EGFR inhibitor Erlotinib across several time points and the corresponding proportion of cells predicted to be arrested/proliferating. **(k)** Principal component analysis illustrating the superimposition of single cell RNA-seq profiles (circles) of G0 arrested NSCLC cells before/after EGFR inhibition onto the bulk RNA-seq reference data (triangles) for MCF10A cells occupying various stress response states. **(l)** The proportion of NSCLC cells in **(k)** predicted to occupy different stress response states across several time points.

**Figure 7:**
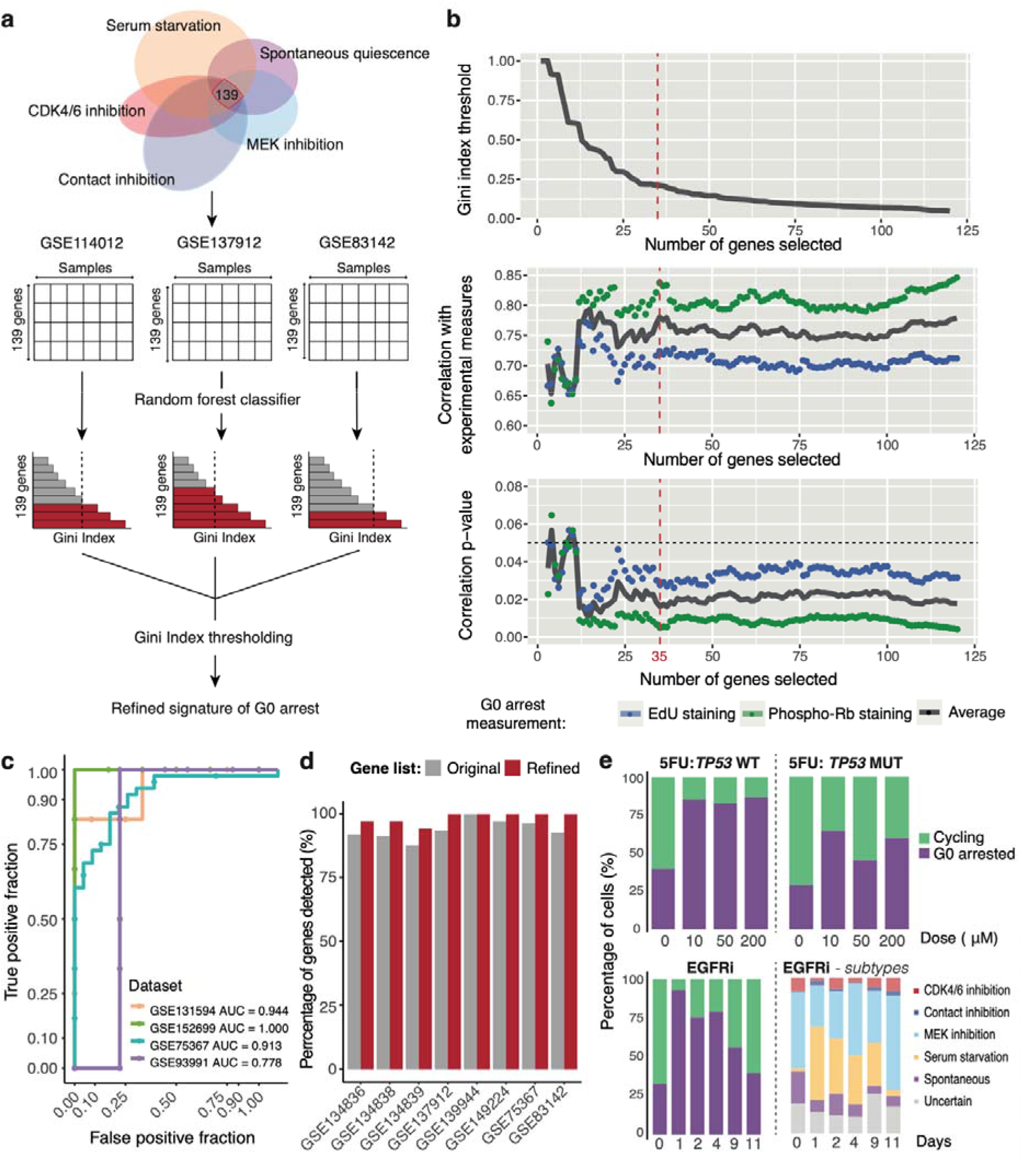
Optimisation of the G0 arrest signature for use in single cell RNA-seq data. **(a)** Methodology for refining the gene signature of G0 arrest: random forest classifiers are trained to distinguish arrested from cycling tumours on three high confidence datasets; Gini index thresholding is optimised to prioritise a final list of 35 genes. **(b)** Gini index variation, correlation with experimentally measured quiescence via EdU and phospho-Rb staining assays, and corresponding p-values are plotted as the number of genes considered in the model is increased. The red dotted line indicates the threshold chosen for the final solution of 35 genes. The black dotted line indicates the threshold for p-value significance. **(c)** Additional external validation of the 35 gene signature acting as a classifier of G0 arrested and proliferating cells in single cell and bulk datasets. **(d)** Dropout in single cell data by gene signature. The percentage of genes out of the 35 (red) and 139 (grey) gene lists with reported expression across the single-cell RNA-seq datasets analysed in this study. **(e)** Proportion of cycling and G0 arrested cells estimated in single cell datasets of p53 wild type and mutant lines treated with 5FU, as well as cells treated with EGFR inhibitors. Data as in Figure 6.

While G0 arrest may confer an overall survival advantage in most cancers, it can also provide a pool of cells that are capable of developing resistance to therapy(12, 99). Using our methodology, we indeed observed an increase in G0 arrest levels in cell lines following treatment with EGFR, BRAF and CDK4/6 inhibitors, as well as conventionally used chemotherapies such as 5- Fluorouracil (5-FU) in multiple bulk RNA-seq datasets (Figure 6d). Furthermore, the recent widespread availability of single-cell transcriptomics offers the opportunity to investigate the impact of G0 arrest on such therapies with much greater granularity than is allowed by bulk data. Using our G0 arrest signature and single-cell data from RKO and SW480 colon cancer cell lines treated with 5-FU(100), we could observe G0 arrest and proliferation decisions following conventional chemotherapy treatment. Within the p53 proficient cell line RKO, the fraction of G0 arrested cells increased from 41% to 93% after treatment with a low dose (10 μM) of 5-FU and persisted at higher doses (Figure 6e-f). In contrast, a comparable increase in G0 arrest was not observed in *TP53* mutant SW480 cells, further emphasizing the key role of p53 as a regulator of cell cycle exit (Figure 6g-h). This implies that although *TP53* mutations confer a more aggressive tumour phenotype and may drive resistance via other mechanisms, *TP53* wild-type tumour cells are more likely to be capable of entering a G0 “persistent” state associated with drug resistance. SW480 cells showed higher apoptotic activity following treatment compared to RKO cells, particularly within actively cycling cells, further corroborating that cells capable of entering G0 arrest may be intrinsically less vulnerable to this therapy (Supplementary Figures 8a- b).

Similarly, using single cell data from an *EGFR* mutant Non-Small Cell Lung Cancer (NSCLC) cell line treated with the EGFR inhibitor Erlotinib(101), we predicted that 40% of cells were likely to exist in a G0 arrest state prior to treatment. EGFR inhibition led to a massive decrease in cell numbers immediately after treatment, mostly due to proliferating cells dying off (Supplementary Figures 8c-d), while the proportion of arrested cells increased to 96% at day 1, indicating an immediate selective advantage for such cells (Figure 6i-j). These cells appear to gradually start proliferating again in the following days during continuous treatment, with the percentage of proliferating cells approaching pre-treatment levels by day 11 (Figure 6j). The same trend captured by our signature could be observed upon KRAS and BRAF inhibition in different cell line models (Supplementary Figure 8e-h, Supplementary Table 3)(12, 101). Furthermore, during the first days of treatment the NSCLC cells that survived EGFR inhibition appeared to reside in a state most resembling that induced by serum starvation (Figures 6k-l). Both EGFR kinase inhibitors and serum starvation have been shown to trigger autophagy(102), which may explain the convergence between this inhibitory trigger and the type of stress response. At day 11 most of the remaining arrested cells appeared in a state similar to that preceding the treatment (Figure 6l).

Thus, G0 arrest appears to explain resistance to broad acting chemotherapy agents as well as targeted molecular inhibitors of the Ras/MAPK signalling pathway, being either selected for, or induced immediately upon treatment, and gradually waning over time as cells start re-entering the cell cycle. Using massively multiplexed chemical transcriptomic data, we also analysed responses to 188 small molecule inhibitors in cell lines at single-cell resolution(103) (Supplementary Figure 9). We observed a large increase in G0 arrest following treatment with compounds targeting cell cycle regulation and tyrosine kinase signalling, consistent with our previous results, but also for compounds modulating epigenetic regulation, e.g. histone deacetylase inhibitors – thus highlighting the broad relevance of G0 arrest.

While links between G0 arrest and therapeutic resistance are prevalently observed in cell lines, one would question whether this translates to similar pathology in cancer patients. While we observed significantly higher G0 arrest levels in pre-treated tumours of non-responders to neoadjuvant chemotherapy in a breast cancer study by Hatzis et al(104) (Supplementary Figure 10a), surveying various targeted therapy datasets from the SELECT study(105) and the TCGA data for links with response to various therapies (single agent and combinations) showed little to no evidence for a bulk signature of G0 being useful for *<u>predicting</u>* resistance in these studies (Supplementary Figures 10b-c). Although the studies available for inspection are rather sparse, evidence from all the analyses presented here suggests there is no universal a priori role that G0 arrest has within the pre-treated primary tumour in determining response to treatment, with more favourable overall outcomes observed occasionally due to slower progressing malignancy, but also resistance observed in the case of chemotherapy in breast cancer. Instead, the role of G0 arrest in enabling therapeutic resistance as a short-lived acquired phenotype as demonstrated in single cell datasets appears more consistent.

### Tumour cell G0 arrest signature for use in single-cell transcriptomics data

Our ability to probe the nature of G0 arrest phenotypes in single cell RNA-seq data using a defined G0 signature could aid the development of methods to selectively target G0 arrested drug-resistant persister cells. However, a major challenge of single cell RNA-seq data analysis is the high percentage of gene dropout, which could impact our ability to evaluate G0 arrest using the full 139 gene signature. The single cell RNA-seq datasets we analysed exhibited an average drop-out of 8.5 genes out of the full gene signature. While our scoring method remains robust to such levels of dropout (Supplementary Figures 1d-f), we also employed machine learning to reduce our initial list of 139 markers of quiescence to a robust 35-gene signature, comprised mainly of RNA metabolism and splicing-regulating factors, but also of genes involved in cell cycle progression, ageing and senescence, which could be applied to sparser datasets with larger levels of gene dropout (Methods, Figure 7a-b, Supplementary Table 4). The optimised signature of G0 arrest performed similarly to the initial broadly defined programme in distinguishing fast cycling tumours from those containing high proportions of G0 cells (Figure 7c), it showed an average dropout of only 0.5 genes across the single cell RNA-seq datasets used in this study (Figure 7d), was similarly prognostic (p=0.004) and showed comparable profiles of resistance to treatment (Figure 7e, Supplementary Figure 11). This minimal expression signature could be employed to track and further study emerging G0 arrest-enabled resistance in a variety of therapeutic scenarios.

## DISCUSSION

Despite its crucial role in cancer progression and resistance to therapies, G0 arrest in all its forms remains poorly characterised due to the scarcity of suitable models and biomarkers for large-scale tracking in the tissue or blood. The lack of proliferative markers such as Ki67 or CDK2(31, 106) does not uniquely distinguish G0 arrest from other cell cycle phases, e.g. G1. Miller et al(32) have shown that the Ki67 is expressed at the mRNA level but the protein is degraded continuously both in G0 and G1, and it rather acts as a graded marker of S/G2/M. Similarly, CDK2 activity is low in G0, G1, builds up at the restriction point, is high in the S phase and is then replaced by CDK1 in mitosis. Reduced CDK2 expression can manifest due to quiescence, mitosis, but also to DNA damage(107), and thus cut-offs to uniquely distinguish its activity in G0 would be difficult to define. Furthermore, these and other reliable markers of G0 arrest such as p27 or p130(50) are best captured at protein level, which is much more sparsely measured, and expression does not accurately reflect their activity. This study overcame this limitation by employing genes active in different forms of quiescence whose patterns of expression are distinct from markers of a longer G1 phase, and capture cell cycle arrest as also observed in senescence, stemness or dormancy. We have extensively validated our method and signature in single cell datasets and cancer cell lines, and have demonstrated that it can reliably and robustly capture signals of G0 arrest both in bulk tissue as well as in single cells.

Within bulk tissue, we are limited in our capacity to distinguish between large fractions of cells residing in short-lived G0 arrest and a smaller fraction of cells that are in deeper G0 arrest, as our score seems to reflect both parameters to a certain extent. Bearing in mind this limitation, our score could potentially also be used in a single cell setting to capture longer-lived cell cycle arrest states such as ones demonstrated in senescence or dormancy, and could assist in identifying such states, but only with the help of additional cell-state specific, immune or secretory biomarkers. Indeed, gene activity linked to cell cycle arrest is not exclusive to quiescence, but can be shared with senescence or dormancy in certain scenarios, as also demonstrated in some of our analyses of senescent cells. This makes it difficult to clearly distinguish states like dormancy, senescence, quiescence (particulary deep quiescence), as even their definitions can be contentious at times both in the context of human cancers (4, 57, 108, 109, 110) as well as in physiological conditions in other organisms (111, 112). Since our signature was derived and validated in experiments that were tailored specifically to induce and/or measure quiescence, we believe the signature proposed in this study best reflects a quiescent-like, reversible G0 cell cycle arrest state. While senescent and dormant cells could be distinguished from their quiescent counterparts simply based on additional senescence and dormancy markers, further research is nevertheless required in the future to delineate signatures that are both necessary and sufficient to unambiguously discriminate all three states. In the meantime, future studies utilising our G0 signature should also test for such additional markers like β-galactosidase activity, the SASP and other senescence markers, or *NRF2*, *NR2F1*, *SOX9*, RARβ(113, 114) and other dormancy markers to ascertain the type of G0 arrest that is being captured.

The versatility of our signature is evidenced by high classification accuracies across a variety of solid cancer datasets. More variable performance was observed when applied to hematopoietic stem cells as it was not designed to capture signals in this context (Supplementary Figure 1b). While we cannot exclude that the patterns captured may also occasionally reflect cell cycle arrest in G1 or G2, this broad signature would still capture phenotypes resulting from intrinsic or extrinsic cellular stress that reflect temporary tumour adaptation during the course of cancer evolution or upon treatment with drugs. Thus, studying such states is relevant for identifying vulnerabilities that could be exploited at different time points during the course of cancer treatment.

We show that G0 arrest is pervasive across different solid cancers and generally associated with more stable, less mutated genomes with intact DNA damage repair pathways. We also find a link between APOBEC mutagenesis and higher levels of G0 arrest. Some neoplastic events enriched in tumours with increased G0 arrest, such as p16 or *ZMYM2* deletions, could mark elevated genomic stress that renders cells more prone to cell cycle exit. We also identified mutational events affecting a variety of genes such as *PTEN*, *CEP89, CYLD*, *LMNA* that appear unfavourable to cell cycle arrest, thus potentially implicating them in influencing G0 arrest-proliferation decisions. Among these, we propose and validate *CEP89* as a novel modulator of G0 arrest capacity in non-small cell lung cancer. A recent paper describes how increased CEP89 copy number and expression correlates with a worse prognosis in ovarian cancer(115), which we hypothesise could be linked to Cep89’s role in modulating G0 arrest. Although we do not yet know how Cep89 regulates G0 arrest, two roles have been ascribed to Cep89 which could be significant. First, Cep89 is required for primary cilium assembly(116, 117). The primary cilium acts as a signalling hub, transducing extracellular signals to intracellular signalling networks, many of which regulate growth and proliferation(118). Cep89 deficiency also leads to defects in Complex IV assembly in the electron transport chain in mitochondria, leading to decreased mitochondrial function and ATP production(119). Decreased ATP would impair the ability of cells to proliferate. Since Cep89 is a coiled-coil protein with no obviously targetable regulatory domains, it will be important to ascertain which Cep89 function is key to regulating the balance between proliferation and arrest in cancer cells to be able to potentially target that process, rather than Cep89 itself, to induce or maintain G0 arrest.

These large-scale genomic associations with G0 arrest phenotypes are only currently feasible in bulk datasets. However, bulk sequenced data has a major limitation in capturing an average signal across all cells within the tumour, which prevents individual cell state identification and counting. Our subsequent exploration of single cell datasets across 193 therapeutic scenarios complements this analysis and illustrates the power of applying our signature in single cells.

Our signature of G0 arrest is prognostic and marks primary tumours with a lower proliferative capacity before treatment, but we also clearly demonstrate that it can be employed to track resistance to multiple cell cycle, kinase signalling and epigenetic targeting regimens, where it often appears as a short-lived phenotype. While this discrepancy may appear incompatible at first glance, it is not unlike other cellular processes that have been shown to present dual roles in a cancer setting, such as reactive oxygen species(120), but also p38α(121) or NRF2(122), both of which have been implicated in quiescence or dormancy(114, 123). It is possible that there is a tipping point between G0 arrest acting beneficially or detrimentally during tumour development and treatment. Furthermore, this is likely influenced by a myriad of other complex factors that we have not had the chance to analyse in depth here, and in some cases it may just be the baseline for acquiring cancer cell stemness or senescent properties. While we acknowledge this conundrum requires further study, we believe this phenotype also offers a unique opportunity to further understand mechanisms of tumour resistance. A key open question remains: if G0 arrest drives resistance, does it do so in a Darwinian fashion, as a pre-existing population that is selected for upon drug treatment, or is it instead an acquired phenotype? Our single cell analyses cannot exclude either scenario. Given the variable links to treatment response and lack of clear evidence for relapse when surveying G0 arrest in primary tumours *before* treatment, it is likely our G0 arrest signature in its current form cannot be used to predict resistance to chemotherapy or targeted therapy, and we would not recommend it for this purpose unless further validated in a specific cancer setting. We have also not inspected the role of G0 arrest in the context of immunotherapy, which remains an area of future study. However, we believe our signature has high value for the study of emerging resistance in an *in vitro*/*in vivo* setting, as a short-lived enabler of drug tolerance. The optimised signature we propose for single cell data makes it tractable to a variety of future studies in this area.

In a treatment setting, vulnerabilities of G0 arrested cells could be exploited for combination therapies. Cells which have exited the cell cycle utilise several mechanisms to achieve drug resistance, including upregulation of stress-induced pathways such as anti-apoptotic BCL-2 signalling(124), anti-ROS programmes(28) or immune evasion(15). Further studies are needed to elucidate which of these mechanisms are specifically employed on a case-by-case basis.

Our findings contribute to the understanding of the aetiology and genetic context of G0 arrest in cancer. This is particularly relevant to identifying new anti-proliferative targets, but also for the detection and eradication of drug tolerant persister cells, which have been frequently, although not always, observed to be slow cycling or entirely quiescent/senescent(8, 9). Importantly, the state of G0 arrest that we have studied here is distinct from that of disseminated tumour cells causing clinical dormancy and cancer relapse, often after many years from the treatment of the primary tumour(4, 125). Here, we have focused on understanding how tumours make proliferation and G0 arrest decisions during the earlier stages of cancer development, within the treatment-naïve primary tumour and as an immediate response to anti-cancer therapies. However, since the dormancy of disseminated tumour cells is fundamentally enabled through a long but temporary cell cycle arrest, we believe our findings of the fundamental processes linked with G0 arrest could in the future help inform a better characterisation of dormant tumour cells when combined with specific microenvironmental signatures that are critical for enabling that process.

## CONCLUSIONS

Overall, our study provides, for the first time, a pan-cancer view of G0 arrest and its evolutionary constraints, underlying novel mutational dependencies which could be exploited in the clinic. We propose a G0 arrest signature which can be robustly measured in bulk tissue or single cells and could potentially inform therapeutic strategies in the longer term. This signature could be assessed in the clinic to track rapidly emerging resistance, e.g. through liquid biopsies or targeted gene panels. We hope these insights can be used as building blocks for future studies into the different regulators of G0 arrest, including epigenetics and microenvironmental interactions, as well as the mechanisms by which it enables therapeutic resistance both in solid and haematological malignancies.

## METHODS

### Selection of G0 arrest marker genes

#### Generic G0 arrest markers

Differential expression analysis results comparing cycling immortalised, non-transformed human epithelial cells and cells in five different forms of quiescence (spontaneous quiescence, contact inhibition, serum starvation, CDK4/6 inhibition and MEK inhibition) were obtained from Min and Spencer(28). A total of 195 genes were differentially expressed in all five forms of quiescence under an adjusted p-value cut-off of 0.05. This gene list, reflective of a generic G0 arrest phenotype, was subjected to the following refinement and filtering steps: (1) selection of genes with a unidirectional change of expression across all five forms of quiescence; (2) removal of genes involved in other cell cycle stages included in the “KEGG_CELL_CYCLE” gene list deposited at MSigDB; (3) removal of genes showing low standard deviation and low levels of expression within the TCGA dataset, or which showed low correlation with the pan-cancer expression of the transcriptional targets of the DREAM complex, the main effector of quiescence, in TCGA. The resulting 139-gene signature is presented in Supplementary Table 1.

#### G0 arrest stress response-specific markers

Gene lists representing spontaneous quiescence, contact inhibition, serum starvation, CDK4/6 inhibition and MEK inhibition programmes were obtained using genes differentially expressed in each individual quiescence form using an adjusted p-value cutoff of 0.05. The gene lists were subjected to filtering steps 2 and 3 described above. Following the refinement steps, 10 upregulated and 10 downregulated genes with highest log2 fold changes were selected for each stress response type.

### Quantification of G0 arrest in tumours

The *GSVA* R package was used to implement the combined z-score(40), ssGSEA(41) and GSVA(42) gene set enrichment methods. For the above three methods a separate score was obtained for genes upregulated in quiescence and genes downregulated in quiescence, following which a final G0 arrest score was obtained by subtracting the two scores. The singscore single sample gene signature scoring method(43) was implemented using the *singscore* R package. In addition to these, we also calculated a mean scaled G0 arrest score based on the refined list of genes upregulated and downregulated in quiescence, as well as a curated housekeeping genes from the “HSIAO_HOUSEKEEPING_GENES” list deposited at MSigDB, as follows:

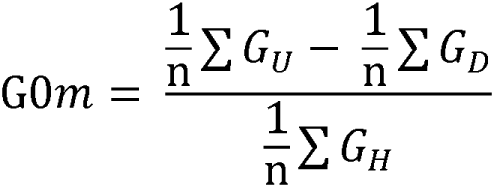

G0*m* = mean scale G0 arrest score

G_U_ = expression of genes upregulated in quiescence

G_D_ = expression of genes downregulated in quiescence G_H_ = expression of housekeeping genes

n = number of genes in each gene set

G0 arrest scores for the TCGA cohort were derived from expression data scaled by tumour purity estimates. The pan-cancer TCGA samples were also classified into groups with “high” or “low” levels of G0 arrest based on k-means clustering (k=2) on the expression data of 139 G0 biomarker genes, following the removal of tissue-specific expression differences using the *ComBat* function from the *sva* R package(126).

### Measuring the duration of G0 arrest

We employed the GSE124109 dataset from Fujimaki et al(52) where rat embryonic fibroblasts were transcriptomically profiled as they moved from short to long-term quiescence in the absence of growth signals. The derived G0 arrest scores using our combined z-score methodology increased from short to longer-term quiescence.

### Validation of G0 arrest scoring methodologies

#### Single-cell RNA-sequencing validation datasets

Datasets were obtained from the ArrayExpress and Gene Expression Omnibus (GEO) databases though the following GEO Series accession numbers: GSE83142, GSE75367, GSE137912, GSE139013, GSE90742 and E-MTAB-4547. Quality control analysis was standardised using the *SingleCellExperiment*(127) and *scater*(128) R packages. Normalisation was performed using the *scran*(129) R package.

#### Bulk RNA-sequencing validation datasets

Datasets were obtained from the GEO database through the following GEO Series accession numbers: GSE93391, GSE114012, GSE131594, GSE152699, GSE124854, GSE135215, GSE99116, GSE124109, GSE61130, GSE64553 and GSE63577. GSE114012 count data were normalised to TPM values using the *GeoTcgaData* R package. All normalised datasets were log-transformed before further analysis.

The accuracy with which the G0 arrest scoring methods could separate proliferating and quiescent samples within the validation datasets was determined by calculating the area under the curve of the receiver operating characteristic (ROC) curves, using the *plotROC* R package.

### Experimental validation in lung adenocarcinoma cell lines

The average fraction of cancer cells spontaneously entering quiescence was estimated for nine lung adenocarcinoma cell lines (NCIH460, A549, NCIH1666, NCIH1944, NCIH1563, NCIH1299, NCIH1650, H358, L23) using EdU and phospho-Rb staining proliferation assays.

Cell lines were obtained from ATCC or Sigma and regularly checked for mycoplasma. A549 and NCIH460 were cultured in DMEM (Gibco). NCIH358, NCIH1299 and NCIH1563 were maintained in RPMI-1640 (Gibco) supplemented with 5mM sodium pyruvate and 0.5% glucose. NCIH1944, NCIH1666, NCIH1650 and L23 were grown in RPMI-1640 ATCC formulation (Gibco). A427 were cultured in EMEM (ATCC). A549, NCIH460, H358, NCIH1299, NCIH1563, A427 were supplemented with 10% heat inactivated FBS. NCIH1666 with 5% heat-inactivated FBS and all other cell lines with 10% non-heat inactivated FBS. All cell lines had penicillin-streptomycin (Gibco) added to 1%. Cells were maintained at 37°C and 5% CO_2_. To calculate the quiescent fraction, A549 and NCIH460 cells were plated at a density of 500 cells/well, and all other cell lines at a density of 1000/well, in 384well CellCarrier Ultra plates (PerkinElmer) in the relevant media. 24h later, 5 μM EdU was added and cells were incubated for a further 24h before fixing in a final concentration of 4% formaldehyde (15 min, RT), permeabilization with PBS/0.5% Triton X-100 (15 min, RT) and blocking with 2% BSA in PBS (60 min, RT). The EdU signal was detected using Click-iT chemistry, according to the manufacturer’s protocol (ThermoFisher). Cells were also labelled for phospho-Ser807/811 Rb (phospho-Rb) using Rabbit mAb 8516 (CST) at 1:2000 in blocking solution, overnight at 4°C. Unbound primary antibody was washed three times in PBS and secondary Alexa-conjugated antibodies were used to detect the signal (ThermoFisher, 1:1000, 1h at RT). Finally nuclei were labelled with Hoechst 33258 (1 μg/ml, 15 min RT) before imaging on a high-content widefield Operetta microscope, 20x N.A. 0.8. Automated image analysis (Harmony, PerkinElmer) was used to segment and quantify nuclear signals in imaged cells. Quiescent cells were defined by the absence of EdU or phospho-Rb staining, determined by quantification of their nuclear expression (Figure 1e-f).

Endogenous PCNA was labelled at the N-terminus with a cDNA encoding mRuby in both A549 and NCI-H1944 cells, using AAV-mediated gene-targeting, according to methods described in Zerjatke et al(54). mRuby-expressing cells were sorted into 50:50 conditioned:fresh media at single-cell density into 96-well plates by FACS and single-cell clones expanded. For live-cell imaging, 500 cells in phenol-red free media were plated per well of a 384 CellCarrierUltra plate (PerkinElmer) the day before imaging. Prior to imaging, a breathable membrane was applied to the plate and cells were imaged on the Operetta HCS microscope (PerkinElmer) at 37 °C, 5% CO2 using the 20x N.A. 0.8 objective and at 10-12 min intervals for 48h. Images were then exported and G0/G1 length (time from mitotic exit to S-phase entry) was analysed manually in FIJI.

The G0 arrest scores for cancer cell lines were calculated using corresponding log-transformed RPKM normalised bulk RNA-seq data from the Cancer Cell Line Encyclopedia (CCLE) database(130).

CEP89 was depleted by ON-Target siRNA Pool from Horizon. NCI-H1299 cells were reverse transfected in 384 well plates with 20nM of Non-targeting control (NTC) or CEP89-targeting siRNA using Lipofectamine RNAiMax (ThermoFisher), according to the manufacturer’s instructions. Cells were left for 24h, before 5 μM EdU was added for the final 24h and then cells were processed as above to determine the quiescent fraction. To determine the level of Cep89 depletion by western blot, cells were reverse transfected with siRNA in 24 well plates. 48h after transfection, cells were lysed directly in 1x SDS sample buffer with 1mM DTT (ThermoFisher). Samples were separated on pre-cast 4-20% Tris-Glycine gels, transferred to PVDF using the iBlot2 system and membranes blocked in blocking buffer (5% milk in TBS) for 1h at RT. The membrane was then cut and the upper half was incubated in 1:1000 Cep89 antibody (Sigma, HPA040056), the bottom half in B-actin antibody 1:2000 (CST; 3700S) diluted in blocking buffer overnight at 4’C. Membranes were washed three times in TBS-0.05% TritonX-100 before being incubated in secondary anti-rabbit (Cep89) or anti-mouse (B-actin) HRP conjugated antibodies (CST 7074P2 and CST 7076P2, respectively) diluted 1:2000 in blocking buffer for 1h at RT. Membranes were washed three times again and signal detected using Clarity ECL solution (BioRad) and scanned on an Amersham ImageQuant 800 analyser.

### Multi-omics discovery cohort

FPKM normalised RNA-sequencing expression data, copy number variation gene-level data, RPPA levels for p27 as well as mutation annotation files aligned against the GRCh38 human reference genome from the Mutect2 pipeline were downloaded using the *TCGABiolinks* R package (131) for 9,712 TCGA primary tumour samples across 31 solid cancer types. Haematological malignancies were excluded as the G0 markers were derived in epithelial cells and might not be equally suited to capture this phenotype in blood. For patients with multiple samples available, one RNA-seq barcode entry was selected for each individual patient resulting in 9,631 total entries. All expression data were log-transformed for downstream analysis. During G0 arrest score calculation, expression data for the primary tumour samples was scaled according to tumour purity estimates reported by Hoadley et al(132) to account for potential confounding cell cycle arrest signals coming from non-tumour cells in the microenvironment. Samples with purity estimates lower than 30% were removed, leaving 8,005 samples for downstream analysis.

The mutation rates of all TCGA primary tumour samples were determined by log-transforming the total number of mutations in each sample divided by the length of the exome capture (38Mb).

*TP53* functional status was assessed based on somatic mutation and copy number alterations as described in Zhang et al(133). *TP53* mutation and copy number for the TCGA tumours were downloaded from cBioPortal (http://www.cbioportal.org). Tumours with *TP53* oncogenic mutations (annotated by OncoKB) and copy-number alterations (GISTIC score ≤ -1) were assigned as *TP53* mutant and CNV loss. Tumours without these *TP53* alterations were assigned as TP53 wild type. The effects of the *TP53* mutation status on G0 arrest were then determined with a linear model approach with the G0 arrest score as a dependent variable and mutational status as an independent variable. The *P* values were FDR-adjusted.

APOBEC mutagenesis enriched samples were determined through pan-cancer clustering of mutational signature contributions as described in Wiecek et al(134). The APOBEC mutagenesis cluster was defined as the cluster with highest mean SBS2 and SBS13 contribution. This was repeated 100 times and only samples which appeared in the APOBEC cluster at least 50 times were counted as being APOBEC enriched.

Aneuploidy scores and whole genome duplication events across TCGA samples were obtained from Taylor et al(135). Microsatellite instability status for uterine corpus endometrial carcinoma, as well as stomach and colon adenocarcinoma samples were obtained from Cortes-Ciriano et al(78). Telomerase enzymatic activity “EXTEND” scores were obtained from Noureen et al(62). Expression-based cancer cell stemness indices were obtained from Malta et al(63). Centrosome amplification transcriptomic signature (CA20) scores were obtained from Almeida et al(89).

### PHATE dimensionality reduction

The *phateR* R package(136) was used to perform the dimensionality reduction with a constant seed for reproducibility. The *ComBat* function from the *sva* R package(137) was used to remove tissue-specific expression patterns from the TCGA RNA-seq data.

### Cancer stem cell division estimates

The mean stem cell division estimates for different cancer types used in this study were obtained from Tomasetti and Vogelstein(55).

### Mutational signature estimation

Mutational signature contributions were inferred as described in Wiecek et al(134).

### Machine learning of G0 arrest-linked features via ensemble elastic net regression models

The COSMIC database was used to source a list of 723 known drivers of tumorigenesis (Tiers 1+2). 285 oncogenes and tumour suppressors from a curated list showed a significant enrichment or depletion of mutations or copy number variants in samples with high levels of G0 arrest either pan-cancer or within individual TCGA studies.

To classify G0 arrest-prone from fast proliferating tumours, the 285 genes were used as input features for an ensemble elastic net regression model along the tumour mutational rate, whole-genome doubling estimates, ploidy, aneuploidy scores and 15 mutational signatures, which showed a significant correlation with G0 arrest scores either pan-cancer or within individual TCGA studies. The *caret* R package was used to build an elastic net regression model 1000 times on the training dataset of 3,753 TCGA primary tumour samples (80% of the total dataset). Only samples with at least 50 mutations were used in the model, for which mutational signatures could be reliably estimated. For each of the 1000 iterations, we randomly selected 90% of the samples from the training dataset to build the model. Only features which were included in all 1000 model iterations were selected for further analysis. To test the performance of our approach, a linear regression model was built using the reduced list of genomic features and their corresponding coefficients averaged across the 1000 elastic net regression model iterations. When applying the resulting linear regression model on the internal validation dataset of 936 samples, we found a strong correlation between the observed and predicted G0 arrest scores (R = 0.73, p < 2.2e-16).

SHAP values for the linear regression model used to predict G0 arrest scores were obtained using the *fastshap* R package.

### Gene enrichment and network analysis

Gene set enrichment analysis was carried out using the *ReactomePA* R package, as well as GeneMania(138) and ConsensusPathDB(139). Interactions between *CEP89* and other cell cycle components were inferred using the list of cell cycle genes provided by cBioPortal and GeneMania to reconstruct the expanded network with direct interactors (*STAG1*, *CCND2*, *STAT3*). Networks were visualised using Cytoscape(140).

### Gene lists

Genes associated with the G1 phase of the cell cycle were obtained from the curated “REACTOME_G1_PHASE “ list deposited at MSigDB. Genes associated with the G1/S and G2/M phases of the cell cycle were obtained from Tirosh et al (51).

Genes associated with apoptosis were obtained from the curated “HALLMARK_APOPTOSIS” list deposited at MSigDB.

Genes associated with the senescence-associated secretory phenotype were obtained from Basisty et al(60). Lists of genes making up the various DNA damage repair pathways were derived from Pearl et al(141).

Genes associated with contact inhibition were obtained from the curated “contact inhibition” gene ontology term. Genes associated with serum starvation were obtained from the curated “REACTOME_CELLULAR_RESPONSE_TO_STARVATION” list deposited at MSigDB. MEK inhibition was assessed based on the activity of the MAPK pathway as determined using an expression signature (MPAS) consisting of 10 downstream MAPK transcripts(92).

### Validation of the genomic constraints of G0 arrest

For elastic net model feature validation, RNA-seq data was downloaded for six cancer studies from cBioPortal(142), along with patient-matched whole-genome, whole-exome and targeted sequencing data. The 6 datasets used comprise breast cancer (SMC(143) and METABRIC(144)), paediatric Wilms’ tumor (TARGET(145)), bladder cancer, prostate adenocarcinoma and sarcoma (MSKCC(146, 147, 148)) studies. The data were processed and analysed in the same manner as the TCGA data. RNA-seq data for 27 MCF7 cell line strains, alongside cell line growth rates and targeted mutational sequencing data were obtained from Ben-David et al(79).

### Genomic dependency modelling in breast cancer

An ANOVA-based feature importance classification was used and identified 30 genomic features most discriminative of samples with lower and higher than average G0 arrest scores. A random forest model was then built using the identified features and correctly classified samples according to their G0 arrest state with a mean accuracy of 74% across five randomly sampled test datasets from the cohort.

### Survival analysis

Multivariate Cox Proportional Hazards analysis was carried out using the *coxph* function from the *survival* R package. The optimal quiescence score cut-off value of 2.95 was determined pan-cancer using the *surv_cutpoint* function. We also used this function to determine optimal cut-offs for individual cancer types, as presented in Figure 6c.

### Treatment response single cell and bulk RNA-seq datasets

Datasets have been obtained from the GEO database through the following GEO Series accession numbers for the cell line experiments: GSE134836, GSE134838, GSE134839, GSE137912, GSE149224, GSE124854, GSE135215, GSE99116, GSE152699, GSE178839, GSE139944, and the following accession numbers for the patient sample datasets: GSE191127, GSE109211, GSE50509, GSE65185, GSE66399, GSE68871, GSE99898 (Supplementary Table 3). Unified treatment response data for TCGA was obtained from Moiso, medRxiv 2021(149). The *umap* R package was used for dimensionality reduction with constant seed for reproducibility.

### Stress response subtype determination

#### TCGA cohort studies

Samples with evidence of cell cycle arrest characterised by a generic G0 score > 0 were further subclassified based on the most likely form of stress response, among CDK4/6 inhibition, contact inhibition, MEK inhibition, spontaneous quiescence or serum starvation, using stress-specific expression signatures. We opted for a conservative approach and classed each sample with a high level of G0 arrest into a specific stress response subtype if the arrest score for the corresponding programme was higher than one standard deviation of the distribution across the TCGA cohort, and if the score was significantly higher than for the remaining programmes when assessed using a Student’s t test. Samples which could not be classified into any of the five stress response states characterised in this study were classified as “uncertain”.

#### Single-cell RNA seq treatment response datasets

The stress response subtype of individual single cells was inferred by mapping such individual cells onto the reference dataset of MCF10A cells reflecting different forms of G0 arrest obtained from Min and Spencer(28). The *ComBat* R package was used to remove the study batch effect between the expression data to be classified and the reference bulk RNA-seq data. PCA dimensionality reduction analysis was then used on the combined datasets using the *prcomp* R function. For each patient sample or single-cell expression data entry, a k-nearest neighbour algorithm classification was performed using the *knn* function from the *class* R package. During the classification the three nearest reference bulk RNA-seq data points were considered, with two nearest neighbours with identical class needed for classification.

### Optimisation of the G0 arrest signature

We investigated if a subset of the 139 G0 phase-related genes could act as a more reliable marker of cell cycle arrest that would bypass dropout issues in single cell data. This was performed in three steps:

(1) *Assessment of individual importance as G0 arrest marker for a given gene* We collected three high confidence single cell expression datasets separating arrested from proliferating cells. A random forest model was trained on each dataset separately to predict the state (G0 arrest/cycling) of a given cell based on the expression levels of the 139 genes in the signature. The Gini indices corresponding to each gene in the model were normalised to a range of values between 0 and 1, which would reflect how important an individual gene was for determining G0 arrest state relative to the other 138 genes. The procedure was repeated 1000 times for each of the three datasets, and the average Gini coefficients across iterations were stored.
(2) *Prioritisation of gene subsets based on cumulative importance in the model* Genes were placed in the candidate subset if their importance metric was above a given threshold in at least one of the datasets. By gradually increasing the threshold from 0 to 1, different gene combinations were produced.
(3) *External validation of candidate subsets* The gene combinations in (2) were tested for their ability to predict G0 arrest. For this, a separate validation dataset was utilized, which contained gene expression levels for the 139 genes in the 10 lung cancer cell lines previously employed for experimental validation, along with the quiescence state of the lines as inferred by phospho-Rb and EdU staining. For each gene subset, a combined z-score of G0 arrest was calculated from the expression levels as described previously. The correlations between this z-score and the two experimental measurements of quiescence were used to establish the ability of a gene combination to predict quiescence. Among the top performing subsets, a 35 gene signature with a mean correlation of 78% between predicted and measured G0 arrest levels in the test data (p=0.016) showed the highest correlation with phospho-Rb measurements capturing short-lived G0 arrest, the more common state observed in single cell treatment datasets. Therefore, this signature was deemed to achieve the best trade-off between gene numbers and signal capture.

The optimised gene signature is provided in Supplementary Table 4.

### Statistical analysis

Groups were compared using a two-sided Student’s t test, Wilcoxon rank-sum test or ANOVA, as appropriate. P-values were adjusted for multiple testing where appropriate using the Benjamini-Hochberg method. Graphs were generated using the ggplot2 and ggpubr R packages.

## DECLARATIONS

### Ethics approval and consent to participate

All data employed in this study comply with ethical regulations, with approval and informed consent for collection and sharing already obtained by the relevant consortia where the data were obtained from (TCGA, METABRI, MSK-IMPACT).

### Consent for publication

Not applicable.

### Availability of data and materials

The results published here are in part based upon data generated by the TCGA Research Network: https://www.cancer.gov/tcga, METABRIC (https://ega-archive.org/studies/EGAS00000000083), MSK-IMPACT (https://www.mskcc.org/msk-impact), or deposited at cBioPortal (https://www.cbioportal.org/) and GEO (https://www.ncbi.nlm.nih.gov/geo/). The specific GEO datasets employed in the analyses are summarised in Supplementary Tables 2 and 4.

All code developed for the purpose of this study can be found at the following repository: https://github.com/secrierlab/CancerG0Arrest

### Authors’ contributions

MS designed the study and supervised the computational analyses. ARB designed and supervised the experimental validation in cell lines. GB supervised the analysis of p53 functional association. AJW developed the quiescence scoring methodology and performed all computational analyses to validate and apply it in bulk and single cell datasets, as well as link it to genomic features. SC performed the experimental validation of G0 arrest prevalence and CEP89 association with G0 arrest in cell lines. DK performed the inference of the minimal signature of G0 arrest applicable in single cell data. MPC performed the random forest modelling and feature selection in breast cancer. LG helped with the gene prioritisation. GMT wrote the code for batch effect correction and for PCA mapping of single cell data on a reference dataset. DHJ performed the APOBEC enrichment classification. PZ and LX performed the G0 arrest comparison of p53 wild type and mutated cancers. MS, AJW, ARB and SC wrote the manuscript, with contributions from all other authors. All authors read and approved the manuscript.

## Acknowledgements

We would like to thank Prof Chris Barnes for the very helpful discussions and input on the findings of the study.

