## Supplementary Material for "Genomic hallmarks and therapeutic implications of G0 cell cycle arrest in cancer"

### SUPPLEMENTARY TABLES

**Supplementary Table 1. The 139-gene signature of G0 arrest.** The up- or downregulation status in G0 arrest is indicated for each gene.

**Supplementary Table 2. Summary of external G0 arrest score validation datasets.**

**Supplementary Table 3. Summary of external cancer cell line treatment response data.**

**Supplementary Table 4. Results of the signature optimisation analysis.** For distinct signatures of different gene length, the Pearson correlations and corresponding p-values when validating against the experimental EdU and Phospho-Rb measurements are shown, along with the average statistics. The optimal gene signature proposed for use in single cell data is highlighted in red.

### SUPPLEMENTARY FIGURES

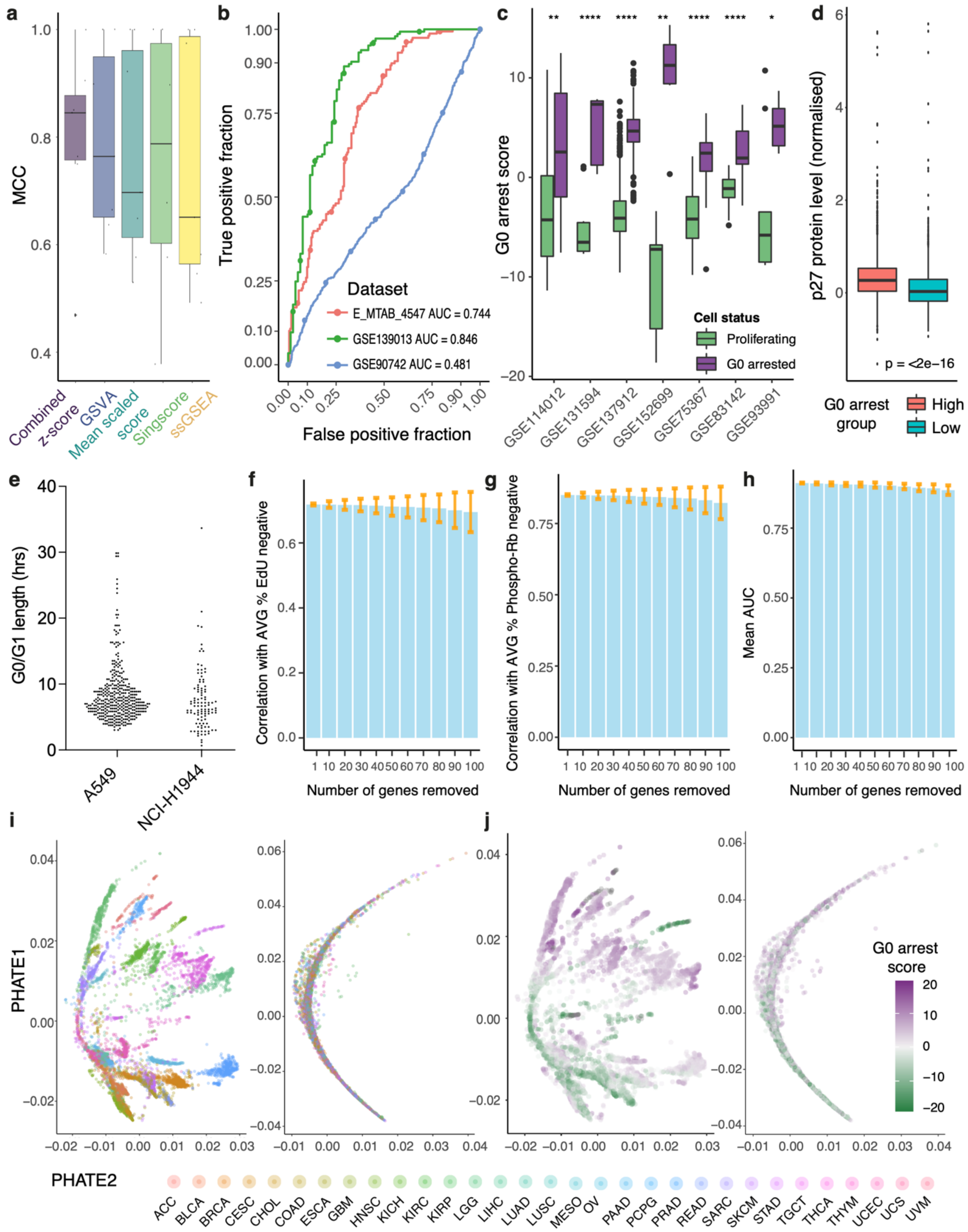


**Supplementary Figure 1: G0 arrest score evaluation and validation. (a)** Classification accuracies measured using the Matthews correlation coefficient (MCC) comparing the combined z-score method to other scoring methodologies for inferring G0 arrest across the seven single-cell and bulk RNA sequencing validation datasets. **(b)** Receiver operating characteristic (ROC) curves illustrating the accuracy of the combined z-score methodology when separating G0 arrested and proliferating hematopoietic stem cells from three publicly available single-cell RNA-seq datasets. Datasets used are denoted by their corresponding GEO series accession number and the area under the curve (AUC) for each dataset is specified. **(c)** G0 arrest scores compared between proliferating and quiescent/dormant single cells individually isolated and sequenced. The x axis indicates the seven single cell validation experiments that the data are derived from. Significant differences are observed in all cases. *p<0.05; **p<0.01; ****p<0.0001. **(d)** Samples with high levels of G0 arrest present significantly higher protein levels of p27. **(e)** G0/G1 length in individual cells over a 48-hour period in two selected lung cancer cell lines. Each dot represents a cell, and the time taken to enter S-phase after mitotic exit is shown on the y axis. **(f)** Mean correlation between computationally inferred G0 arrest scores after randomly removing variable number of genes from the G0 arrest signature and the fraction of cells entering G0 arrest in nine lung adenocarcinoma cell lines, as assessed through an EdU staining assay. **(g)** Same as **(f)** but with experimental levels of G0 arrest measured using a phospho-Rb staining assay. **(h)** Mean classification accuracy across 7 bulk and single-cell RNA-seq validation datasets after randomly removing variable number of genes from the G0 arrest signature. **(i-j)** PHATE plots displaying the first two principal components obtained from dimensionality reduction analysis across 8,005 primary tumour TCGA samples, based on the expression of 139 genes differentially expressed across different G0 arrest programmes before (left) and after (right) removal of tissue specific expression patterns. Each primary tumour sample is coloured by their corresponding tissue of origin **(i)** or the corresponding G0 arrest score **(j)**.

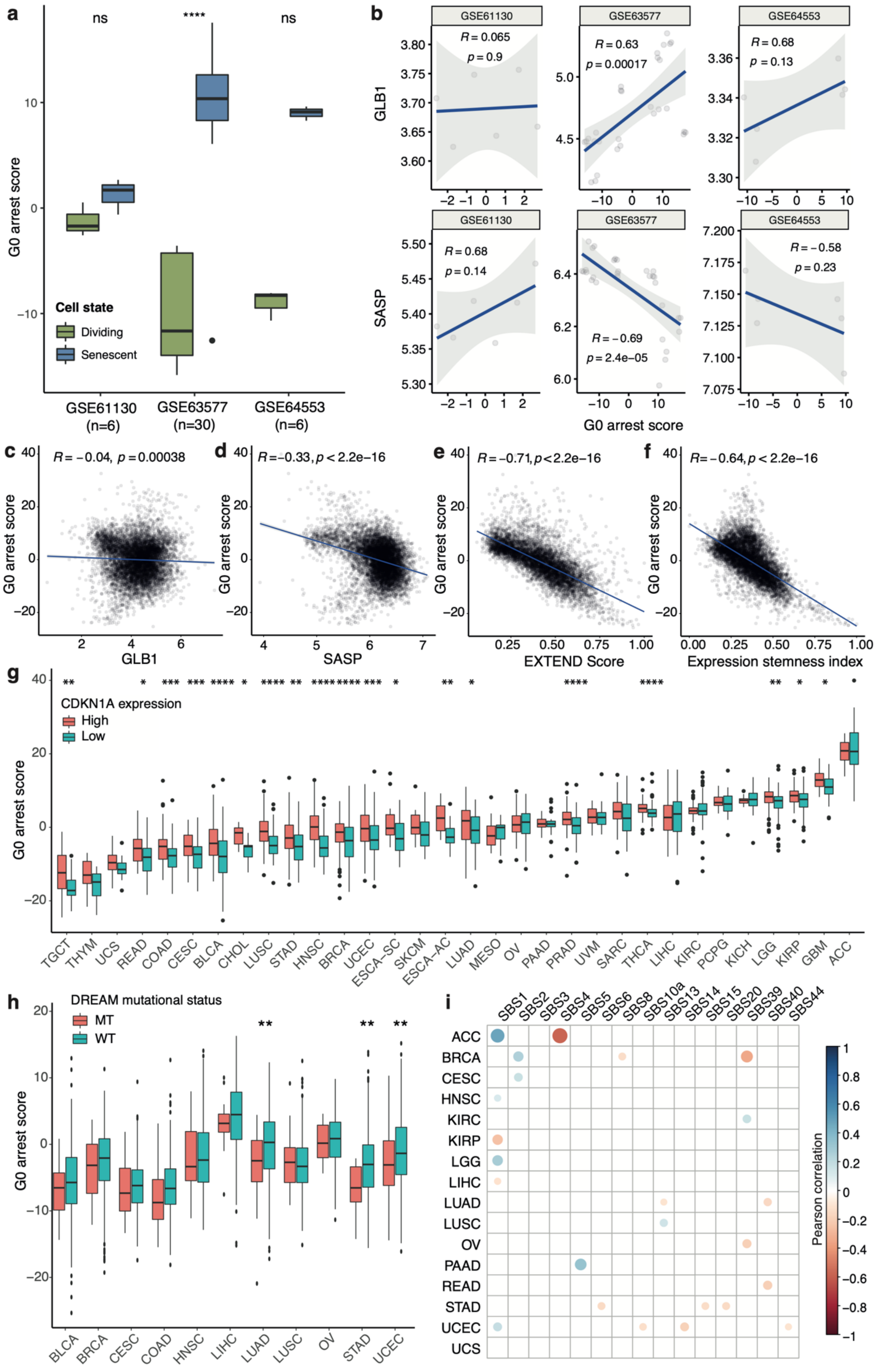


**Supplementary Figure 2: Expression and genomic patterns of G0 arrest in tumours. (a)** G0 arrest scores compared between individually separated dividing and senescent cells from three single cell sequencing experiments. Sample numbers are indicated in brackets. **(b)** Correlation between G0 arrest levels and the classical markers of senescence, β galactosidase (GLB1) and the Senescence-Associated Secretory Phenotype (SASP), in the three single cell senescence datasets tested. Dots correspond to individual cells. **(c-f)** Correlation between G0 arrest scores and **(c)** the expression of the β galactosidase-encoding gene GLB1, **(d)** the expression of the SASP, **(e)** telomerase activity inferred via the "EXTEND" score, and **(f)** the stemness expression index. **(g)** Consistently higher levels of G0 arrest are detected in samples with high CDKN1A expression (encoding p21, upper quartile of the expression distribution) compared to samples with low expression (lower quartile of the expression distribution) across TCGA cancer types. **(h)** Consistently higher levels of G0 arrest are detected in samples with a functional DREAM complex. TCGA studies where mutations within one of the DREAM components could be detected in at least 10 patients are shown. **(i)** Correlation between G0 arrest levels and mutational signature contributions across TCGA cancer types. Only significant correlations are displayed (Pearson correlation p-value $< 0.05$). *$p<0.05$; **$p<0.01$; ***$p<0.001$; ****$p<0.0001$.

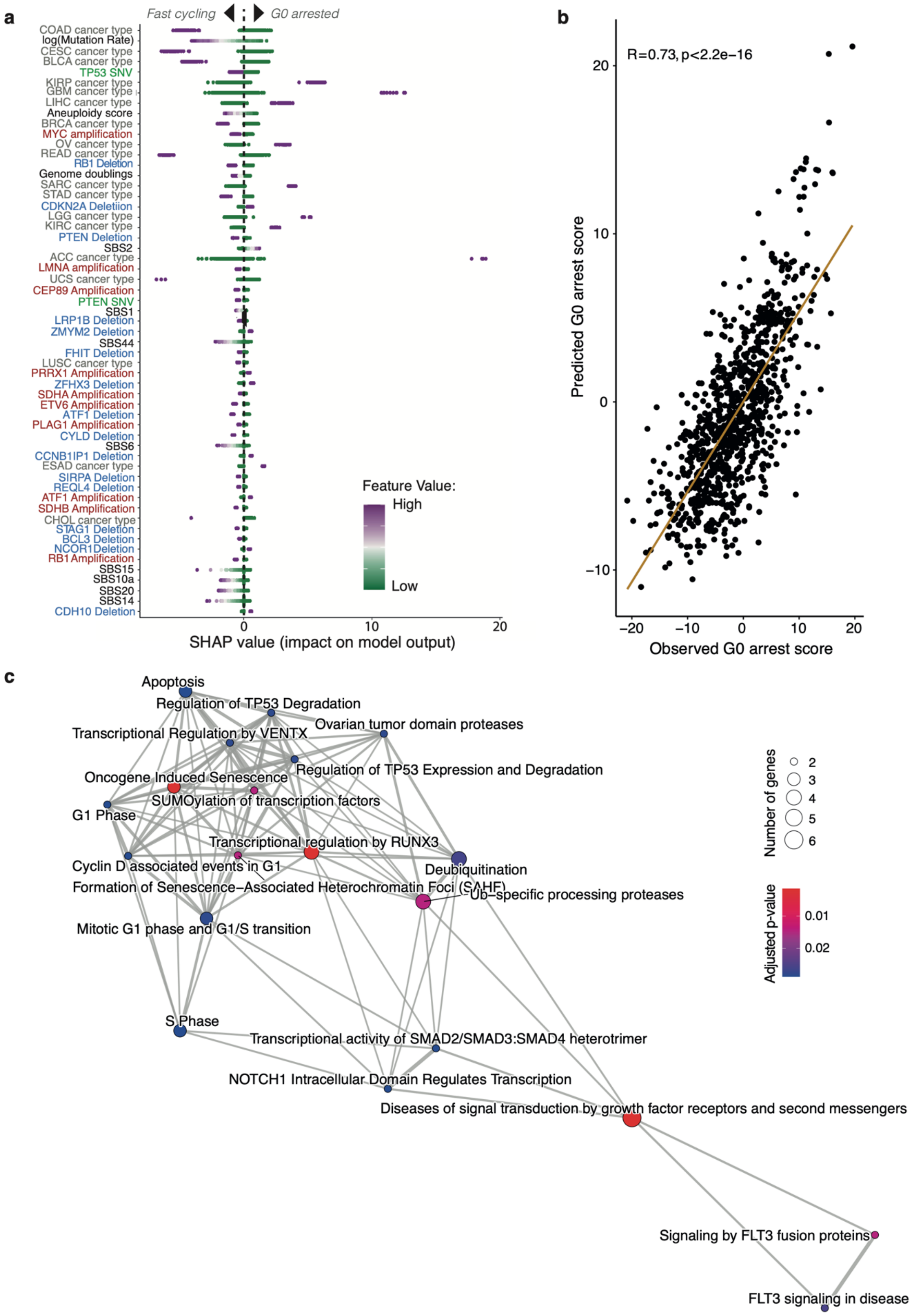


**Supplementary Figure 3: Modelling and validating a pan-cancer classifier of G0 arrest.**
**(a)** Genomic features and cancer types significantly associated with G0 arrest, ranked

according to their importance in the pan-cancer linear regression model (from highest to lowest contribution). The genomic features highlighted are identical to Figure 3c, but cancer types are shown here as well. Each point corresponds to one sample and is coloured according to the original value of the respective feature. For discrete variables purple indicates the presence of the feature and green indicates the absence of the feature. For cancer type variables, purple indicates the patient belongs to the specified TCGA cancer type group. The Shapley values indicate whether the G0 arrest scores predicted by the ensemble elastic net regression model for each patient are increased or decreased depending on the cancer type of each individual as well as specific SNVS, CNV and other genomic features. **(b)** Correlation between the observed G0 arrest scores within the test dataset (x axis) and scores predicted by the linear regression model using the selected pan-cancer features (y axis). **(c)** Pathways significantly altered in association with G0 arrest in tumours.

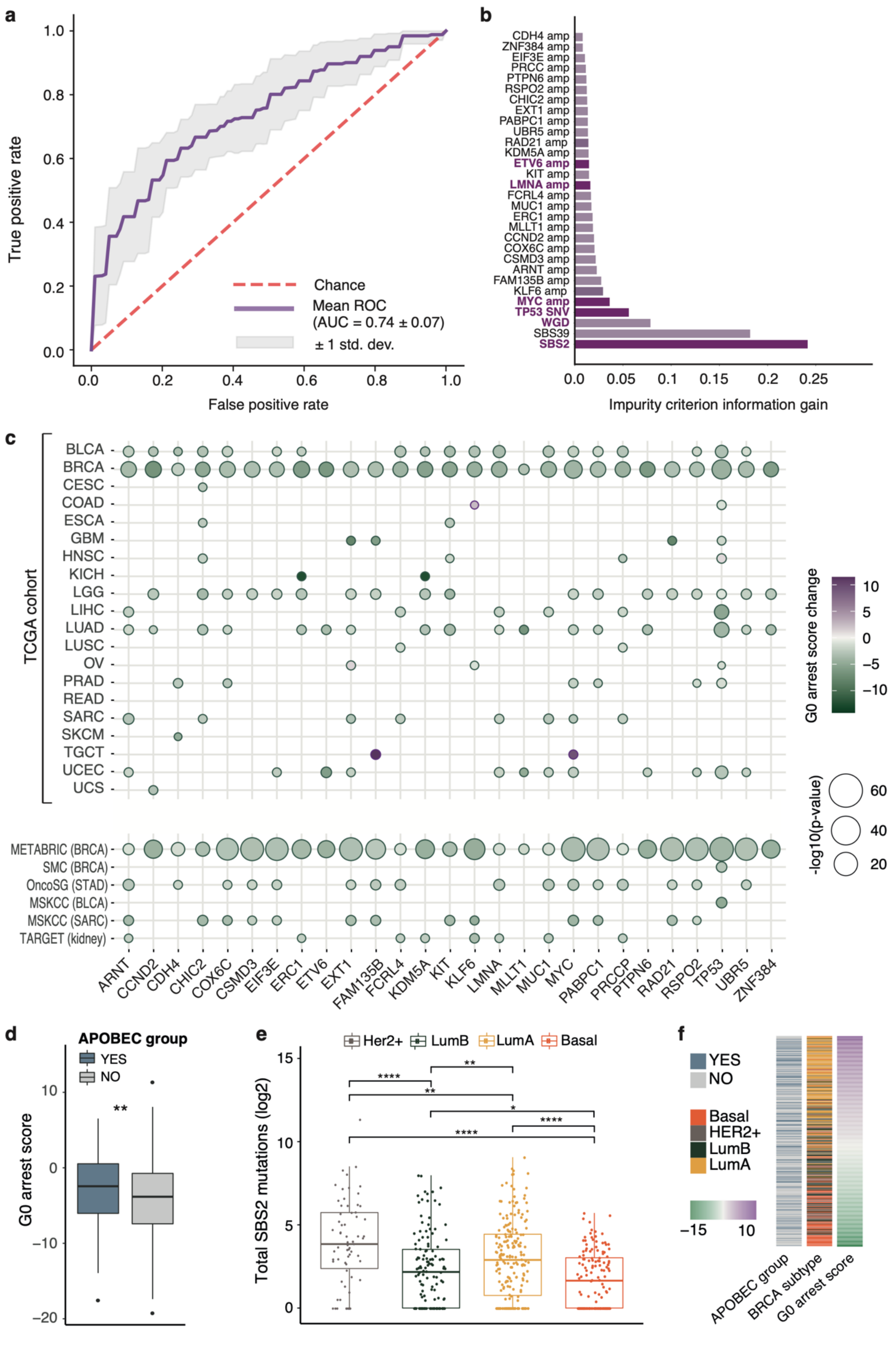


**Supplementary Figure 4: Genomic landscape of G0 arrest in breast cancer. (a)** ROC curve illustrating the mean performance of the random forest model for G0 arrest prediction in breast cancer based on 9 randomisations. **(b)** Impurity criterion information gain for the top ranked genomic events predictive of G0 arrest in the TCGA BRCA cohort, modelled using random forests. Features also appearing in the pan-cancer model are highlighted in dark purple. WGD = whole-genome doubling. **(c)** Tissue-specific changes in G0 arrest estimates between samples with and without G0-associated amplifications (red) and SNVs (purple), identified by the breast cancer specific random forest model, within the TCGA cohort (top panel) and external validation datasets (bottom panel). **(d)** G0 arrest scores are significantly higher in APOBEC mutation-enriched breast cancer samples. **(e)** APOBEC-linked SBS2 mutational burden compared between breast cancer subtypes. **(f)** Association between G0 arrest scores, APOBEC enrichment and breast cancer subtypes in TCGA. *p<0.05; **p<0.01; ****p<0.0001.

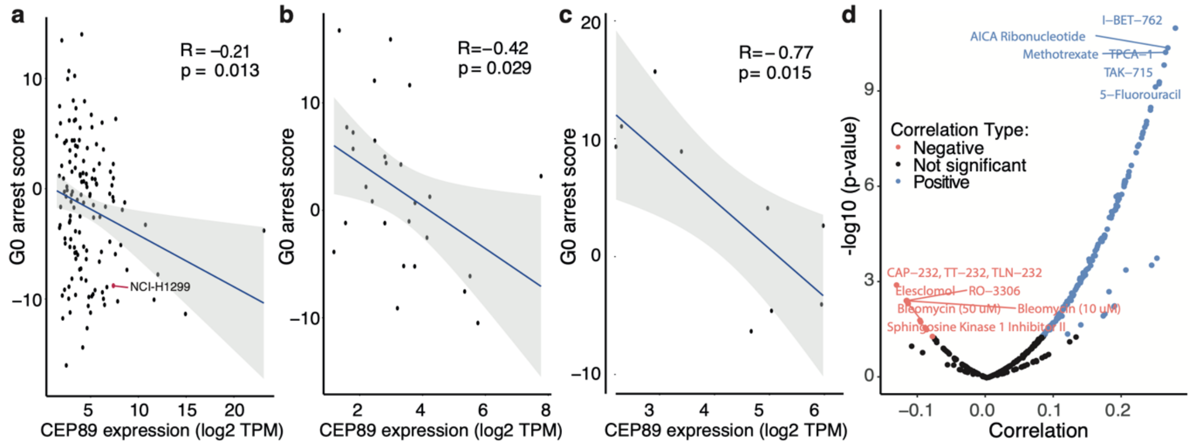


**Supplementary Figure 5: CEP89 expression is associated with G0 arrest and has prognostic value. (a-c)** CEP89 expression is negatively correlated with G0 arrest scores in lung **(a)**, pancreas **(b)** and thyroid **(c)** cancer cell lines. The NCI-H1299 cell line where CEP89 activity was tested is highlighted in red. **(d)** Compounds with selective toxicity on cell lines with high CEP89 expression. The volcano plot shows the Pearson correlation coefficients and the corresponding p-values for associations between CEP89 expression and Area Under the dose-response Curve (AUC) for each compound across cell lines form the CCLE.

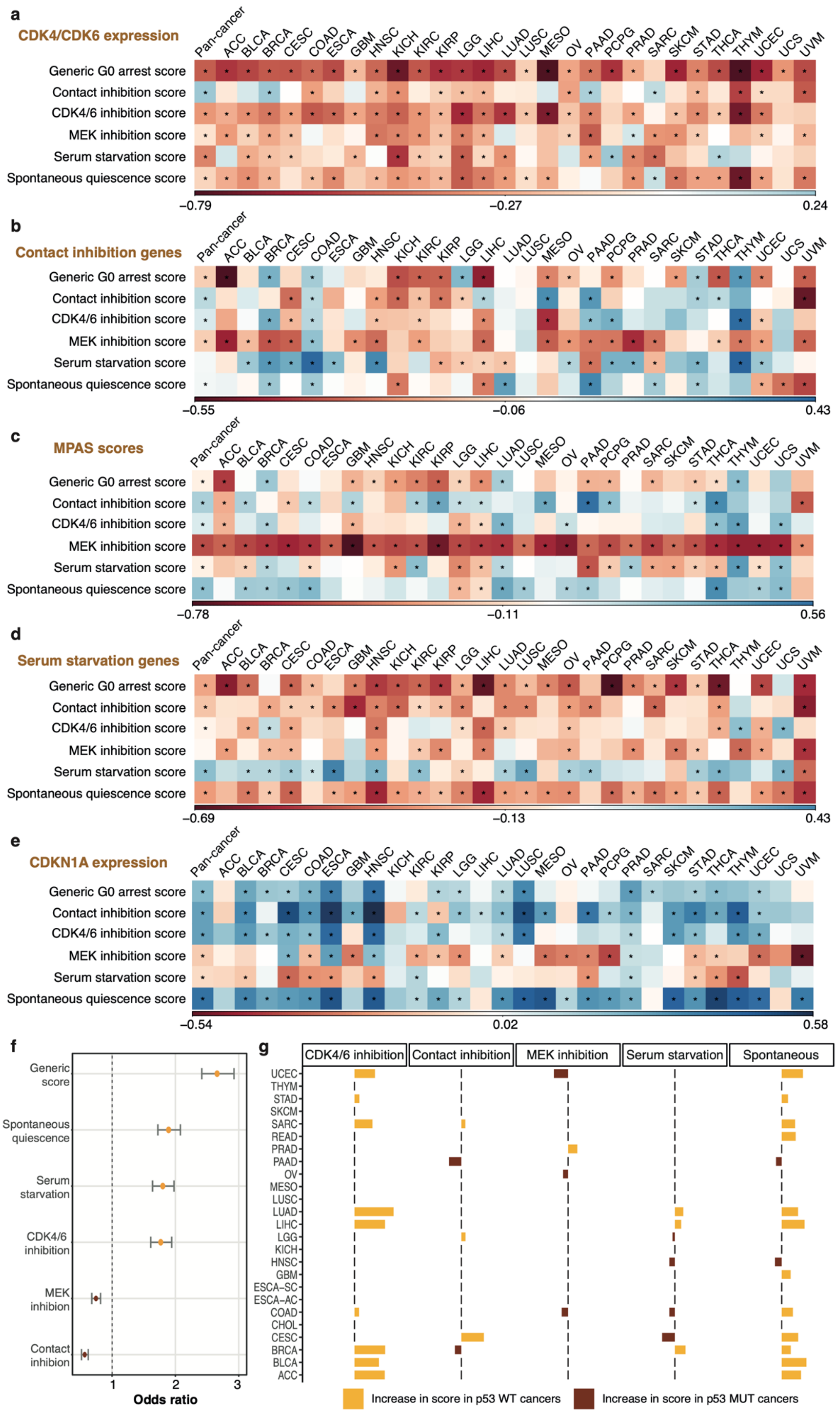


**Supplementary Figure 6: Stress response programme validation and links with p53 status.** Correlation between stress response-specific G0 estimates and **(a)** mean expression of

CDK4/6, **(b)** the mean expression of curated contact inhibition genes, **(c)** MPAS scores, **(d)** mean expression of curated serum starvation genes and **(e)** CDKN1A expression, across individual TCGA cancer studies and pan-cancer. The colour gradient indicates the strength of the association (positive or negative). Asterisks indicate significant associations. **(f)** Enrichment of p53 proficiency in cancers categorised as G0 arrested based on a dominant stress response programme. Enrichment means (yellow) and depletion means (i.e. corresponding to p53 mutant enrichment, brown) are depicted for tumours displaying a specific G0 arrest programme, alongside confidence intervals, across all TCGA cancers. **(g)** Changes in G0 arrest between p53 proficient (WT) and deficient (MUT) cancers, stratified by stress response and cancer study. The mean increase in WT/MUT cancers is depicted for each programme, only when the differences are significant.

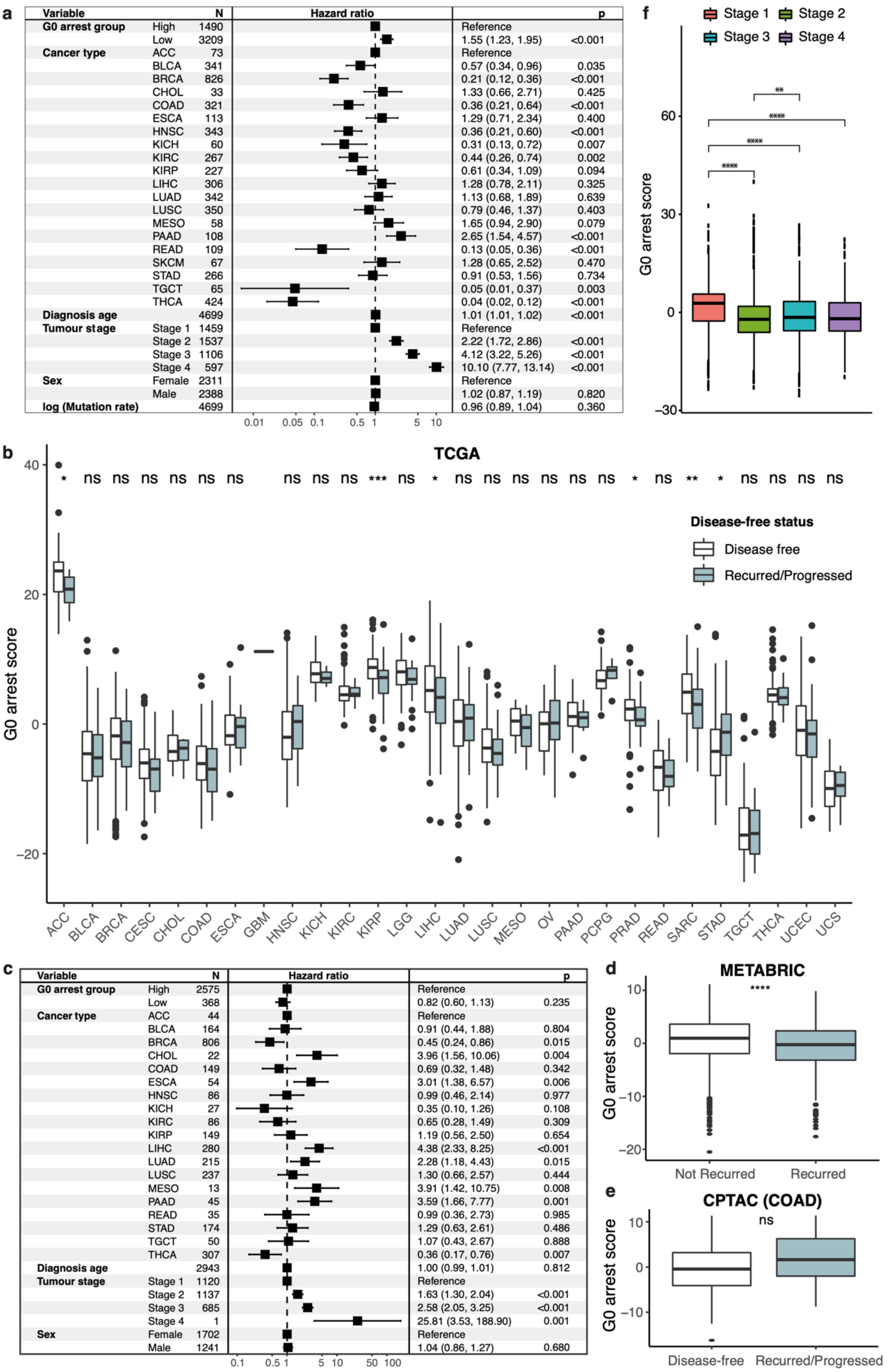

**Supplementary Figure 7: Relevance of G0 arrest to clinical outcome in cancer. (a)** Multivariate Cox proportional hazards analysis of the disease specific survival in the TCGA cohort, modelled on G0 arrest capacity (binarized by optimised survival cut-off into high, where score > 2.95; and low, where score <= 2.95) and potential cofounding factors. **(b)** G0 arrest scores in TCGA cancers compared across tumour stages. **(c)** Comparison of G0 arrest levels in primary tumours based on disease-free or recurrence/progression status across TCGA cancers. **(d)** Multivariate Cox proportional hazards analysis for disease-free survival in TCGA, modelled on G0 arrest capacity and potential confounding factors. **(e)** Comparison of G0 arrest levels in primary tumours based on recurrence in the METABRIC cohort. **(f)** Comparison of G0 arrest levels in primary tumours based on recurrence/progression status using protein-level data from CPTAC in colon adenocarcinoma (COAD). ns = non-significant; * p<0.05; ** p<0.01; ***p<0.001; ****p<0.0001.

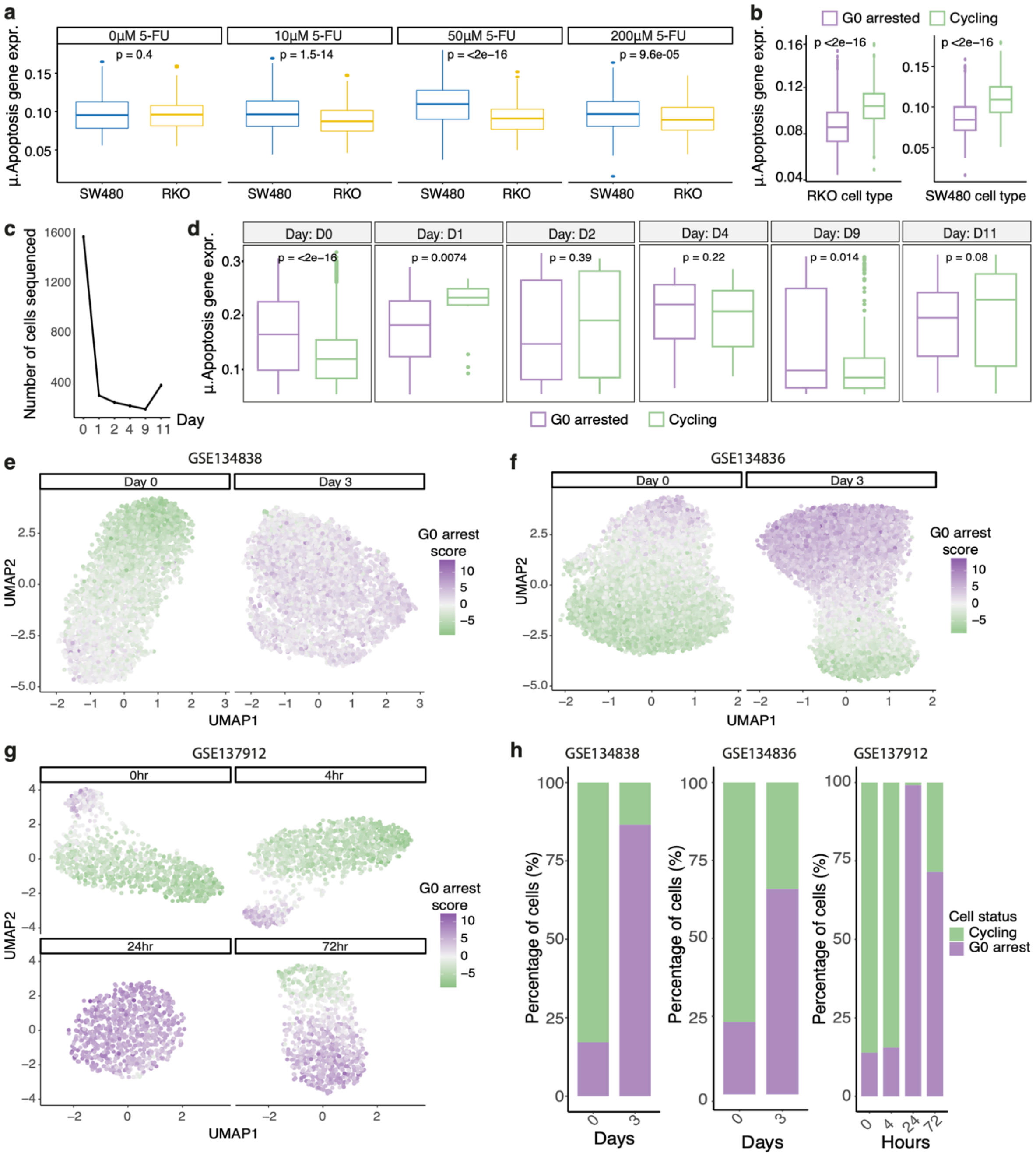

**Supplementary Figure 8: G0 arrest signatures of drug tolerance in single cell data. (a)** The average expression of apoptosis genes compared between TP53 wild type RKO and TP53 mutant SW480 colorectal cancer cell lines in response to various 5-FU treatment doses. **(b)** The average expression of apoptosis genes compared between G0 arrested and cycling cells within the TP53 WT RKO and TP53 MT SW480 cell lines. **(c)** The number of NSCLC cells sequenced during the EGFR inhibition treatment time course. **(d)** The average expression of apoptosis genes compared between G0 arrested and cycling cells at different time points during EGFR inhibition treatment. There is a notable increase in apoptotic activity of cycling cells at day 1. **(e)** Response of melanoma cancer cells to BRAF inhibition treatment across several time points, illustrated using UMAP plots coloured by the G0 arrest score of individual cells. **(f)** Response of NSCLC cells to EGFR inhibition via Erlotinib across several time points, illustrated using UMAP plots coloured by the G0 arrest score of individual cells. **(g)** Response of lung adenocarcinoma cell lines to KRAS inhibition treatment across several time points, illustrated using UMAP plots coloured by the G0 arrest score of individual cells. **(h)** Percentage of cells illustrated in **(e-g)** predicted to be arrested in G0 across the treatment time course.

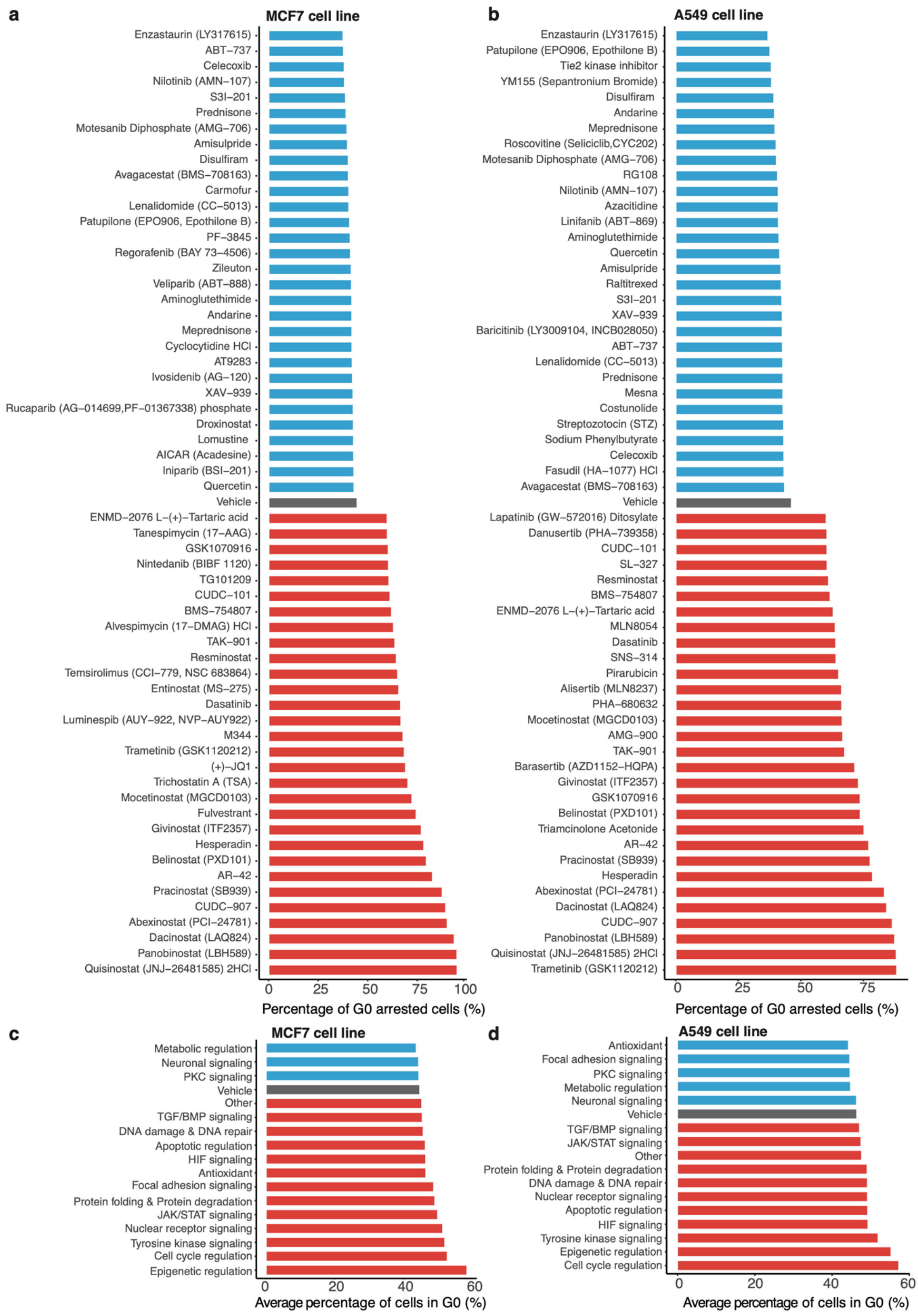


**Supplementary Figure 9: G0 arrest dynamics upon various treatment modalities. (a-b)** Percentage of MCF7 **(a)** and A549 **(b)** cells arrested in G0 following 24-hour treatment with selected compounds. Treatments highlighted in blue and red indicate the 30 compounds which showed the highest decrease and increase in G0 arrest levels, respectively, in comparison to the control, shown in grey. **(c-d)** The average percentage of MCF7 **(c)** and A549 **(d)** cells arrested in G0 following a 24-hour treatment with compounds affecting specific pathways. Pathways shown in blue and red indicate an average decrease and increase in G0 arrest respectively in comparison to the control, shown in grey.

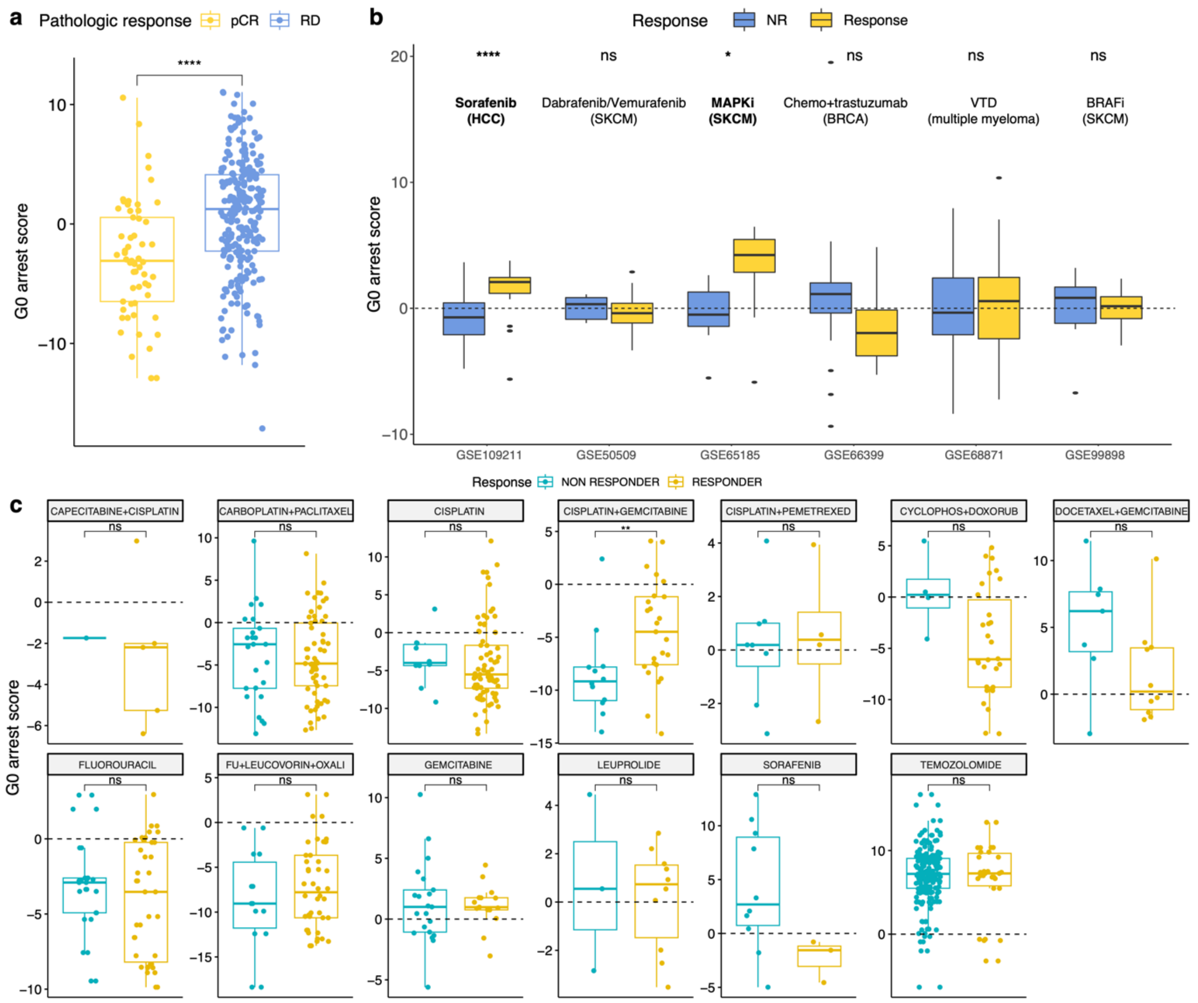


**Supplementary Figure 10: G0 arrest levels in patient samples compared between responders and non-responders to various cancer treatments. (a)** G0 arrest levels of pre-treatment samples taken from breast cancer patients displaying pathologic complete response (pCR) versus residual disease (RD) after neoadjuvant chemotherapy (study GSE25055). **(b)** G0 arrest levels compared between responders and non-responders (NR) to a variety of targeted therapies analysed in the SELECT study. The therapies, cancer type and GEO series are indicated in the plot. **(c)** Differences in pre-treatment G0 arrest levels between responders and non-responders to various first line therapies documented across TCGA cancers. The horizontal dotted line marks the 0 cut-off for what would be considered a tumour with high levels of G0 arrest versus a highly proliferative tumour.

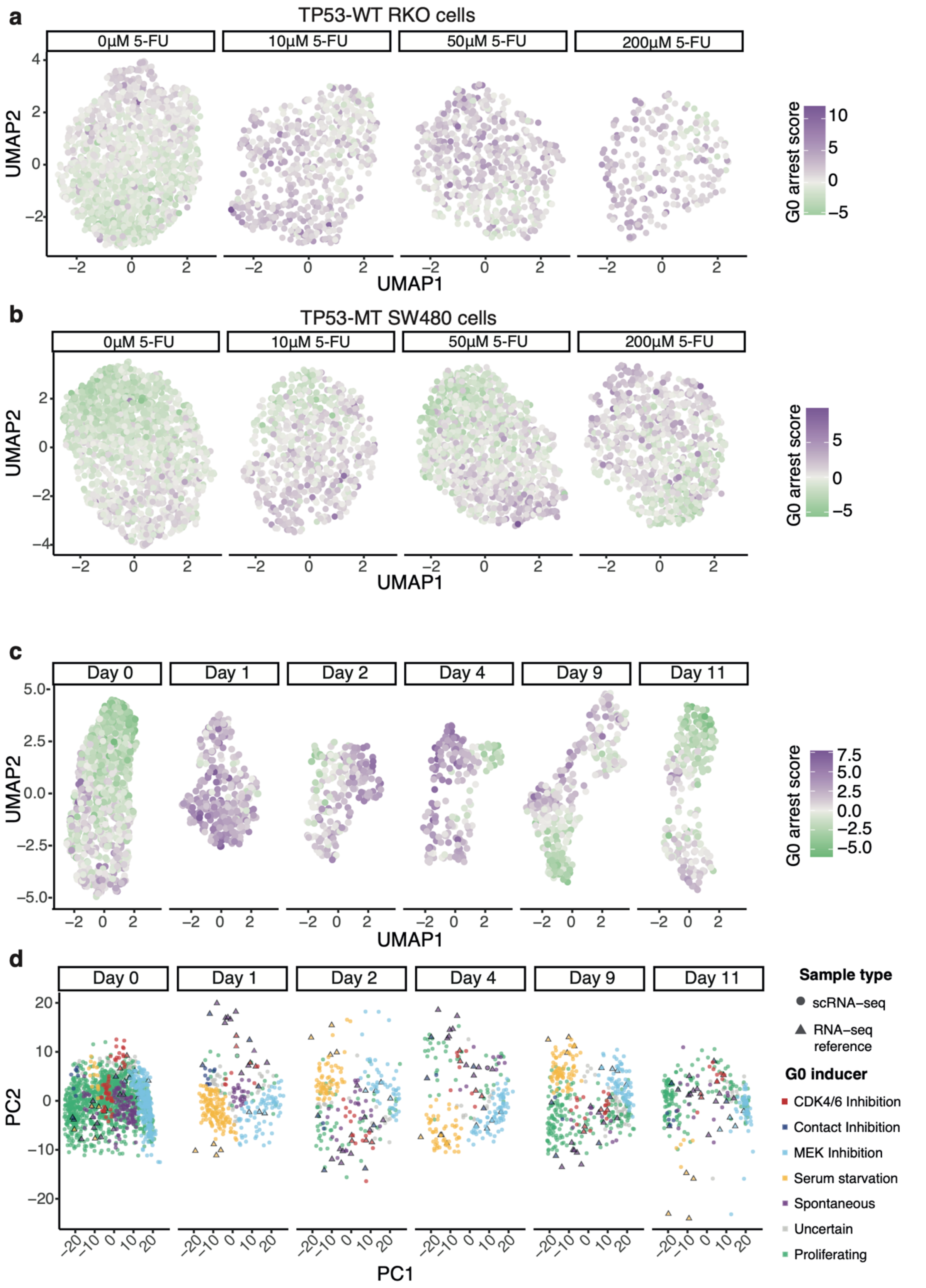


**Supplementary Figure 11: Application of the reduced G0 arrest signature to scRNA-seq data. (a-c)** UMAP plot illustrating the response of **(a)** the TP53-proficient RKO colorectal cancer cell line to various 5-FU doses, **(b)** the TP53-deficient SW480 colorectal cancer cell line to various 5-FU doses, and **(c)** the P9 NSCLC cell line to the EGFR inhibitor Erlotinib across several timepoints. Each dot is an individual cell and it is coloured according to its G0 arrest level, calculated using the optimised 35 gene signature. **(d)** Principal component analysis illustrating the superimposition of scRNA-seq profiles (circles) of G0 arrested NSCLC cells, inferred using the optimised 35 gene signature, before and after EGFR inhibition onto the bulk RNA-seq reference data (triangles) for MCF10A cells occupying various G0 arrest states. The NSCLC cell line stress response programme was predicted using a k-nearest neighbour algorithm based on the bulk RNA-seq reference.
